# High-Resolution, Large Imaging Volume, and Multi-View Single Objective Light-Sheet Microscopy

**DOI:** 10.1101/2020.09.22.309229

**Authors:** Bin Yang, Merlin Lange, Alfred Millett-Sikking, Ahmet Can Solak, Shruthi Vijay Kumar, Wanpeng Wang, Hirofumi Kobayashi, Matthew N. McCarroll, Lachlan W. Whitehead, Reto P. Fiolka, Thomas B. Kornberg, Andrew G. York, Loic A. Royer

## Abstract

Recent developments in Oblique Plane Microscopy (OPM) have shown that it can achieve high spatio-temporal resolution. Here we describe a single objective light-sheet microscope based on oblique plane illumination that achieves: (i) large field of view and high-resolution imaging via a custom remote focusing objective; (ii) fast volumetric imaging by means of *light-sheet stabilised stage scanning* – a novel scanning modality that extends the imaging volume without compromising imaging speed nor quality; (iii) multi-view imaging by alternating the orientation of light-sheet illumination and detection to improve the image quality on large samples; (iv) simpler design and ergonomics by remote placement of coverslips to allow inverted imaging, enabling imaging across scales in a high-throughput format. Overall, we achieved a resolution of 450 nm laterally and 2 μm axially and a field of view of 3000 μm × 800 μm × 300 μm. We demonstrate the speed, field of view, resolution and versatility of our novel instrument by imaging various systems, including zebrafish whole brain activity, *Drosophila* egg chamber development, and zebrafish development – up to nine embryos simultaneously.

## 1 Introduction

In the past 15 years, light-sheet microscopy has become an essential imaging method for biology^1,2^. It has been particularly impactful in developmental biology allowing the first *in-toto* volumetric reconstructions of embryonic development of model organisms such as *Drosophila*, zebrafish, and mouse^1–3^. The ability to image whole developmental arcs, and to follow hundreds of thousands of cells in space and time has shown the potential of light-sheet microscopy in answering long standing questions^3,4^.

However, a major limitation in light-sheet microscopy is the requirement of complex multi-objective configurations that complicate sample mounting. Recent developments have attempted to improve the sample mounting ergonomics with open-top imaging systems^5–7^, or by alleviating the necessity for orthogonal objectives^8^. Among these methods, Oblique Plane Microscope (OPM)^9^ uses a single objective lens (referred to as the primary objective) for both illumination and detection, without additional reflecting elements. The fluorescence is relayed to a secondary objective and a tertiary objective and focused onto a camera. Both the secondary and tertiary objectives are mounted remotely, leaving only the primary objective at the close proximity of the sample. This configuration makes the optical system effectively a “single” objective light-sheet microscope and is compatible with diverse specimens including the intact rodent brain^10^ and multi-well plated samples^11,12^. As a drawback, OPM traditionally was unable to utilize the full numerical aperture (NA) of the primary objective for fluorescence detection, lowering its resolution and sensitivity. This has been addressed more recently via refractive-index-mismatched remote focusing^12–14^, which unlocks the potential of OPM for high resolution imaging by giving it access to the full numerical aperture for detection. However, full NA detection imaging has only been demonstrated with high magnification systems (100x and 60x), for a limited field of view up to 200 μm^14^. This field of view cannot accommodate large living samples such as developing embryos and fundamentally limits imaging throughput. The standard solution to this problem is to tile the acquisition over multiple fields-of-view, sacrificing imaging speed.

Another limitation of light-sheet microscopy applied to large living specimens such as zebrafish embryos is the presence, within the sample, of structures that absorb, refract and scatter, both the illumination and detection light. By illuminating and/or detecting from different orientations, multiview imaging is an effective solution where an image with better coverage and overall better quality can be computationally reconstructed^1,15–18^. However, multi-view imaging often requires either complex multi-objective configurations that complicate sample mounting, or sample rotation that leads to decreased imaging speed.

To address both of these problems, we present DaXi (Da and Xi are both words from Chinese Mandarin, where Da refers to the large imaging volume and Xi for the high resolution capability), a novel single-objective design that is capable of imaging large samples, with a large imaging volume, uncompromised image resolution and speed, and multi-view imaging. DaXi (see Fig. 1, Supplementary Fig 1 and Note 1-2) achieves this goal by means of several innovations: First, we use a novel custom tertiary objective which maintains the image resolution and field of view of a low magnification primary objective (20x). Second, we introduce a new fast 3D scanning modality – *light-sheet stabilised scanning* (LS^3^) – that further extends the effective imaging volume (i.e. scanning range) without compromising imaging speed nor quality. Third, we show how to achieve multi-view imaging and enhance volumetric coverage and image quality with dual illumination and detection. Fourth, we further improve sample mounting ergonomics by converting our microscope from upright to inverted using remote focusing. Last, we demonstrate these novel capabilities by imaging of zebrafish tail development, *Drosophila* egg chamber development, parallel imaging of multiple embryos, and fast recording of whole brain activity in zebrafish larvae.

**Figure 1:**
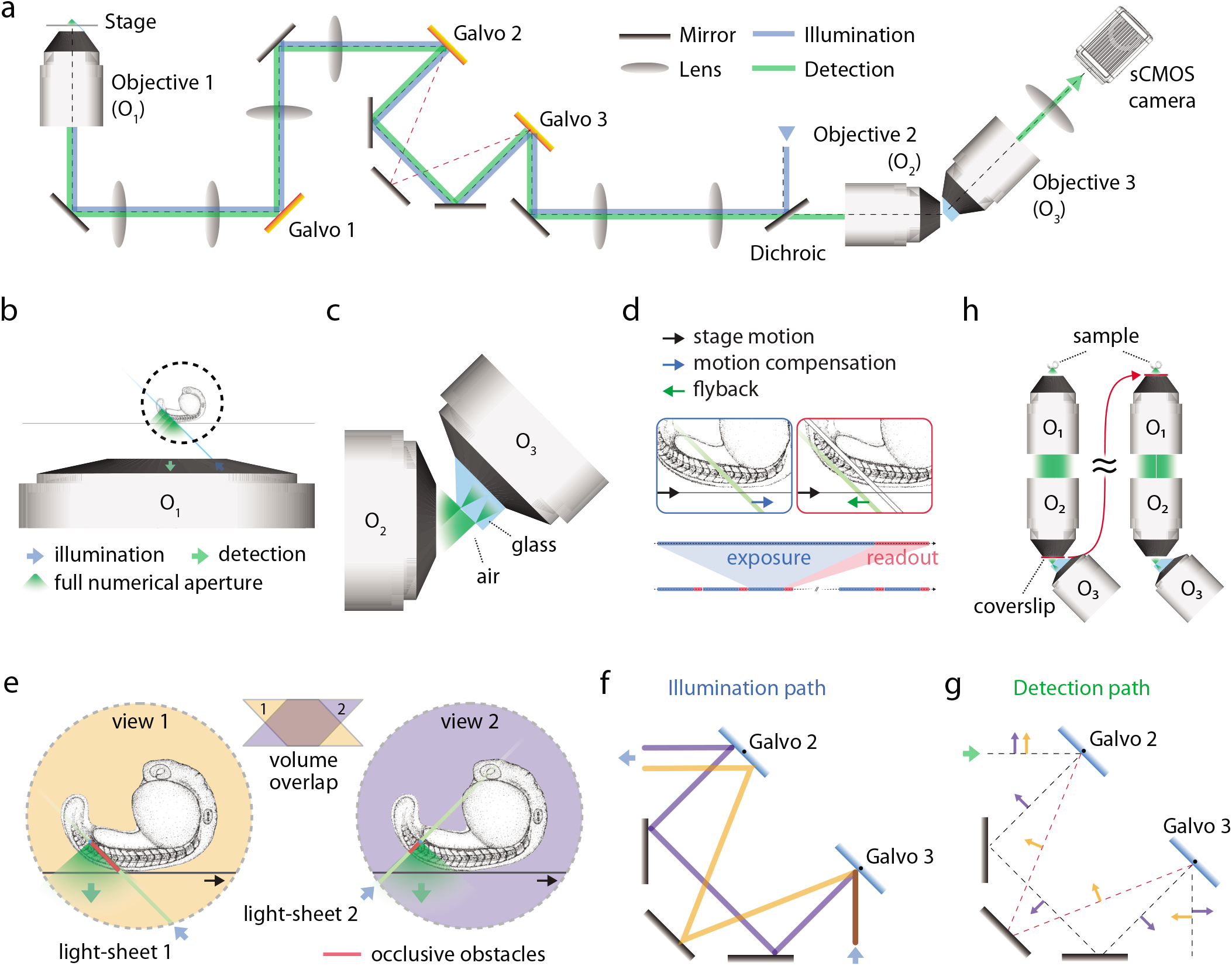
Design of a high-resolution, large field of view, and multi-view single objective light-sheet microscope. **(a)** Simplified scheme of the optical setup. **(b)** In this setup, the light-sheet excitation and emission pass through a single objective. The fluorescence is collected by *O*_1_ and relayed downstream with full NA detection, ensuring high resolution imaging. **(c)** The full NA detection is achieved by oblique remote focusing using a bespoke objective with a monolithic glass tip and zero working distance. The glass tip compresses the collection half-angle allowing a tilt range from 0 to 55°. **(d)** During imaging, the stage moves the sample along the scanning axis. To avoid motion blur, the galvo mirror moves the light-sheet alongside the stage movement during the camera exposure for each image. The galvo mirror flies back during the readout time and restarts this compensatory movement during the exposure of the next frame. This is nearly equivalent to having a stage that is as fast as a galvometer scanner and allows for motion-blur-less volumetric imaging over millimeters. Importantly, illumination and detection planes remain centered along the entire optical train to give optimal light collection, minimal aberrations, and thus pristine image quality. **(e)** Our instrument is capable of dual light-sheet excitation. This improves illumination coverage and image contrast, as for most points within the sample, one of the two light-sheet orientations will have a shorter penetration depth through the sample giving a more contrasted and complete image. The dual view imaging is achieved through an imaging flipping module consisting of two galvo mirrors and three regular mirrors along the optical path (**f** and **g**). **(f)** The illumination light goes along the path highlighted in orange or blue, resulting in opposing incident angle at the sample space. **(g)** Similarly, the fluorescence light goes through either of the two paths, resulting the flipping of the image with respect to that of the other path (blue and orange arrows before and after propagation through the unit), ensuring that the intermediate image is always formed on the front surface of *O*_3_. **(h)** The microscope is converted from upright (dipping, left side) to inverted (immersion, right side) by repositioning the coverslip from the focal space of *O*_2_ to that of *O*_1_, without sacrificing the optical performance.

## 2 Results

### Larger field of view and high resolution by means of a custom objective

Our first goal when designing our microscope was to achieve high resolution over a large field of view. In OPM, a single objective lens (referred to as the primary objective) is used for both illumination (oblique) and fluorescence collection (Fig. 1b). The fluorescence is relayed to a secondary objective with aberration-free remote focusing^19^. A tertiary objective is arranged at the same angle as the illumination lightsheet tilting angle with respect to the secondary objective, to focus the fluorescent signal from the obliquely illuminated imaging plane onto a camera (Fig. 1c). The secondary and tertiary objectives are both mounted remotely, leaving only the primary objective at the close proximity of the sample. This configuration makes the optical system effectively a ‘single’ objective light-sheet microscope, thus facilitating sample mounting and handling.

To maintain the spatial resolution and the field of view of the upstream optical system, the tertiary objective needs to have high NA (at least 1.0), adequate field of view, and reasonable mechanical dimensions to avoid collision with the secondary objective. Previously, both water-immersion^12^ and glass-tiped^13,14^ objectives have been demonstrated to fulfill such requirements for high magnification systems (100x and 60x) with a field of view up to 200 μm. To extend the field of view for low magnification systems (20x in this work), we used a custom tertiary objective (AMS-AGY v2.0) with an NA of 1.0 and a field of view up to 750 μm. This custom tertiary objective is the second instance of a family of objectives that are specifically designed for single-objective light-sheet microscopy (see Fig. 1c, Supplementary Note 3 and Fig 2-4). It features an air-glass imaging boundary and a zero working distance for maximum mechanical clearance and hemispherical collection in air, thus allowing our optical system to achieve uncompromised high resolution with a large field of view.

**Figure 2:**
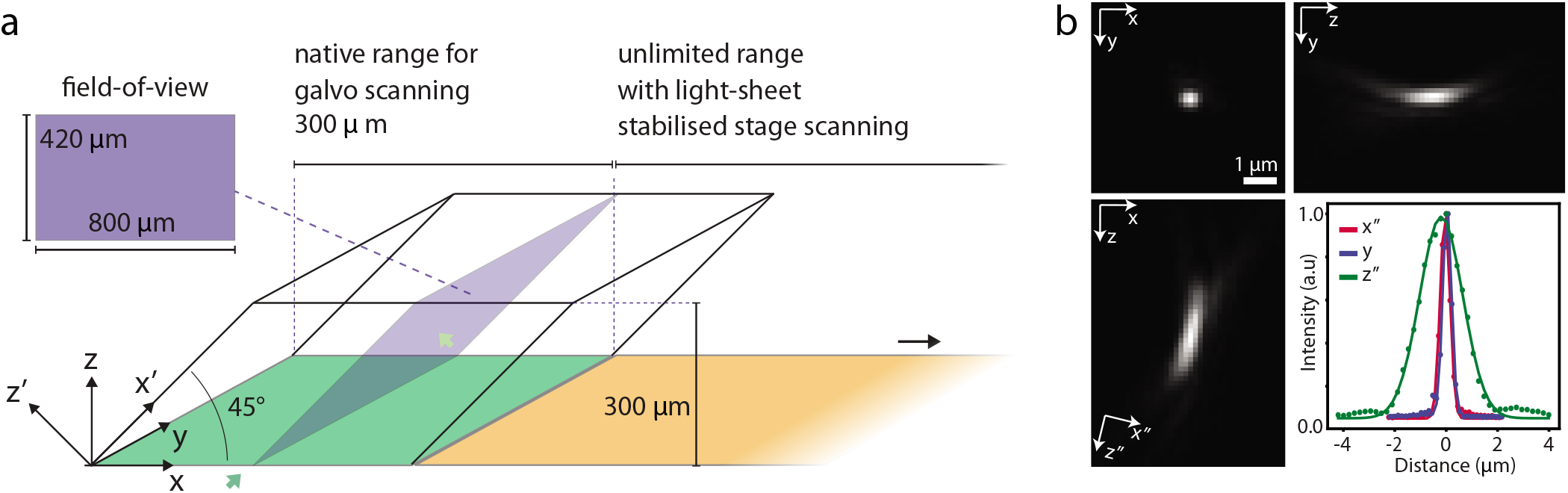
Characterisation of the microscope. **(a)** Imaging volume geometry. The optical axis of the microscope is along the *z*-axis. The coverslip is parallel to the *xy* plane. The sample is illuminated by an oblique light-sheet, in the *x*′*y* plane. The scanning is along the *x* direction. By using light-sheet stabilised stage scanning, the scanning range (up to 75 mm, compared to 300 μm with galvo scanning) is only limited by the stage. **(b)** Point spread function obtained by imaging 100 nm green fluorescence beads. Projections along *xy, xz*, and *zy*, centered line profiles along the three principal axes *x”, y, and z”*, and the FWHMs are respectively 444.1 nm, 403.7 nm and 1984.2 nm.

### Light-sheet stabilised stage scanning

In light-sheet microscopy, a large imaging volume not only requires a large field of view but also a long scanning range as the sample is imaged one plane at a time. Our next challenge was the choice of scan method. Volumetric scanning in a single objective lightsheet microscope is done either by moving the imaging plane (both the illumination and detection) or the sample relative to the other. Moving the imaging plane is faster because it can be implemented with a fast actuator such as a galvo scanner. However, the volumetric scanning range is limited by the field of view of the primary objective. Moving the sample relative to the imaging plane using a microscope stage does not suffer from this and offers a scanning range of more than 10 mm. However, stage scanning either occurs by stepwise motions to avoid motion blur which limits temporal resolution, or by strobing the illumination light, reducing but not completely eliminating motion blur, at the cost of increased peak illumination intensity. More importantly, due to OPM’s scan geometry, stage scanning introduces both axial and lateral blur. This is problematic as the lateral resolution is typically much better than the axial resolution and hence more sensitive to blur. To solve this problem, we developed LS^3^, which combines the high speed of a galvo scanner and the large scan range of a microscope stage (see Fig. 1d and Supplementary Fig. 5). The stage moves continuously while a compensatory motion of the imaging plane induced by a galvo scanner cancels out any relative motion between the sample and the imaging plane during the exposure of one camera frame. The imaging plane is then brought back quickly to the starting position during the camera readout and before a new frame is acquired. All advantages of stage scanning are retained, while simultaneously benefiting from the advantages of high-speed galvo-based scanning. In light of the typical exposures and travel speeds needed in practice, the only true limiting factors of the scanning speed are camera speed and fluorophore brightness.

### Dual light-sheet illumination and detection for multi-view imaging

With the large imaging volume achieved, we are ready to image large samples. The next challenge we faced was that large samples have occluding, refracting, and scattering structures which could reduce image quality. In order to improve the optical coverage and to have consistent image quality, we optically alternate the light-sheet illumination and the viewing direction (orthogonal to the light-sheet) between +/− 45° with respect to the optical axis and imaged the sample sequentially to give a pair of complementary orthogonal views (see Fig. 1e-g). This is achieved by using two galvo mirrors to quickly alternate between two optical paths. One path has two additional mirrors, while the other path adds only one (see Fig. 1f-g, orange and purple paths), hence effectively flipping the light with respect to the optical axis. Importantly, this image flipping module affects both illumination and detection plane simultaneously so that the imaging plane always falls on the glass surface of the tertiary objective regardless of whether the illumination light goes through the sample from left or right. Consequently, both views share the same downstream and upstream optical path, largely simplifying the optical setup and avoiding additional cost. The orthogonal image planes will benefit from the contrasting trajectories into the sample, in many cases avoiding obstacles and therefore returning complementary information that can be fused in post processing.

### Remote coverslip converts a microscope from upright to inverted

Our last consideration was to ensure ease and flexibility of sample mounting, which is often restricted because objectives are either configured for immersion or dipping states. For example, in our instrument, the primary objective (Olympus XLUMPLFLN20XW) does not require a coverslip, whereas the secondary objective (Olympus UPLXAPO20X) does (left side of Fig. 1h). This configuration is suitable for upright imaging where the primary objective can be dipped into the imaging medium directly without needing to image through a glass coverslip. However, we discovered that the coverslip needed by the secondary objective can be moved to support the sample (right side of Fig. 1h), effectively turning this system into an inverted configuration. Moreover, this was achieved without affecting the image quality according to simulations and experimental results (see Supplementary Fig. 6-7). Our lens configuration considerably simplifies microscope design and increases its versatility: inverted imaging enables many imaging modalities, including multi-well plates and microfluidic devices, as well as on-stage sample manipulation.

### Instrument characterisation

We first measured the optical performance of our instrument. The imaging volume is multiple millimeters × 800 μm × 300 μm (x, y, z, respectively, see Fig. 2a). This is achieved by scanning the stage with the light-sheet stabilised relative to the sample to avoid motion blur. Scanning using a galvo has a limited range of about 300 μm, beyond which the illumination starts to be cropped and most importantly the imaging performance starts to degrade. The custom tertiary objective we use guarantees high spatial resolution by full NA detection; thus, the nominal NA of the microscope is close to that of the primary objective O_1_, i.e. 1.0. In practice, we note that the light is compressed towards one edge of the pupil of the tertiary objective O_3_ along the *x*′-axis due to the air-glass interface between O_2_ and O_3_ (see Supplementary Fig. 8-9). As a result, the effective pupil function is no longer symmetric with respective to the optical axis along the *x*′-axis. We measured the point-spread-function (PSF) using 100 nm green fluorescence beads in both views. The full width half maximal (FWHM) values of the PSF along the three principle axes are 442.0 ± 10.0 nm, 395.2 ± 7.8 nm and 1894.1 ± 141.6 nm respectively (Fig. 2b). Some aberrations remain in the optical system that mostly originate from the primary objective. This affects the PSF especially under oblique remote focusing. Future designs that incorporate a deformable mirror could potentially correct system aberrations and further improve the PSF.

Our instrument achieves a combination of large imaging volume and spatial resolution, demonstrated by imaging of whole zebrafish larvae (Fig. 3a-b and Supplementary Movie 1) and *Drosophila* fly egg chambers (Fig. 3c-d and Supplementary Movie 2). Importantly, these images are obtained from single 3D stack acquisitions and do not require stitching, corresponding to an imaging volume of 3000 μm × 800 μm × 300 μm. The dimension along the scanning direction (x-axis) is only limited by the scanning range of the stage (in our case 75 mm). LS^3^ is key to acquiring such large volumes at high-quality. However, when imaging smaller samples (limited to about 300 μm), galvo scanning is faster because its settling time after one volumetric scan is just a few milliseconds, compared to hundreds of milliseconds for the stage. Importantly, our instrument is capable of both scanning modes, providing flexibility dependent on the desired volume dimensions. The imaging speed largely depends on the scanning range, but also on signal level, scanning step size, etc. For an imaging volume of 2000 μm × 800 μm × 300 μm, we typically achieve an imaging speed of about 10 to 30 seconds per volume. By tuning imaging parameters according to experimental requirements, much higher speeds are achievable, similarly to other single objective lightsheet implementations^12,20^. For instance, we imaged zebrafish larvae whole brain activity using genetically encoded calcium indicators, at 3.3 Hz volumetric imaging rate (see Supplementary Fig. 10).

**Figure 3:**
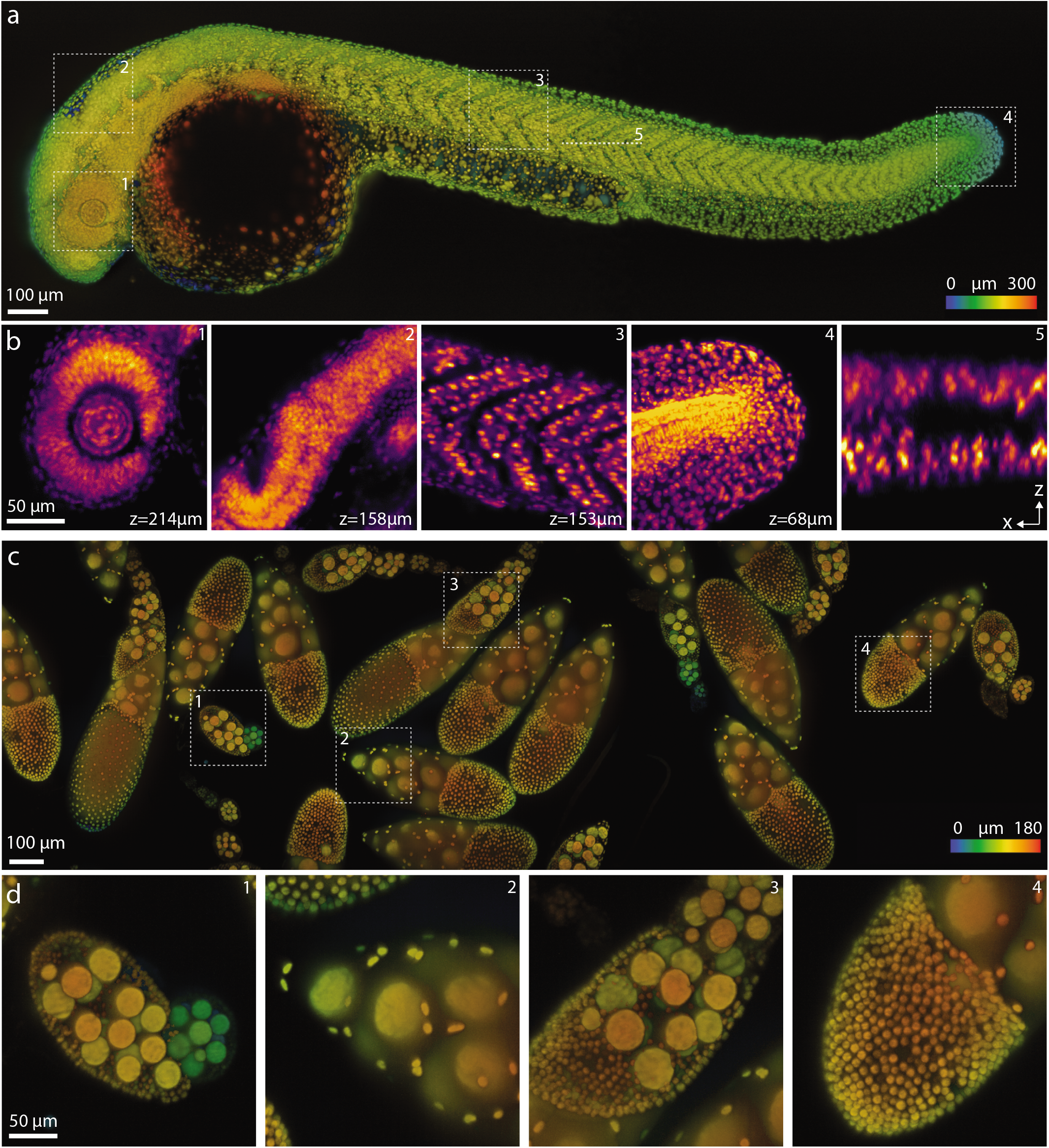
Large volume imaging of *Danio rerio* larval development and *Drosophila melanogaster* egg chambers. **(a)** Images of a zebrafish larvae (~ 30hpf, nuclei labelled with *tg(h2afva:h2afva-mCherry)* imaged using the microscope. Imaging volume (x,y,z) is 3000 μm×800 μm×300 μm acquired every 50 seconds (two views). The depth is color-coded, where blue and red indicate respectively close to and far from the coverglass. **(b)** Four xy slices from different regions (1-4, dashed squares in **a**) at various depths and one xz slice (5, dashed line in **a**) are highlighted. **(c)** Images of *Drosophia* fly egg chambers. Nuclei of germline cells (large) and somatic cells (small) were labeled by expressing UAS-NLS-GFP under the control of Usp10-Gal4 (BDSC-76169). Imaging volume is 3000 μm× 800 μm× 180 μm acquired every 30 seconds (single views) for 3 hours. The depth is color-coded as above. **(d)** Fours regions (dashed squares in **c**) are highlighted.

### Imaging zebrafish tail development

Together, the high resolution, large imaging volume and multi-view features of Daxi enabled us to follow the zebrafish tail development at high spatio-temporal resolution over long periods of time. For example, we imaged the tail extension of a 24 hpf (hours post fertilisation) embryo for 8 hours. To minimize mechanical stress around the animals and allow their normal development, embryos are embedded in exceptionally soft (0.1%) agarose gel in a glass-bottom Petri dish. Sample holding stability during imaging mainly comes from the gravity of the sample itself as well as agarose viscosity. Fig. 4a-b shows that the whole tail can be imaged from the posterior part of the tail (the tail bud) to the anterior trunk, spanning more than a millimeter of tissue along the anterior-posterior axis. The two views obtained by dual orthogonal illumination maximised coverage and image quality. Indeed, as shown in Fig. 4c these views are complementary in terms of contrast and coverage. Depending on the region inspected, one view, or the other, will be better. In general, we find a good agreement between our observations and our assumption that given a point within the sample, the longer the light-sheet penetration depth, the poorer the image quality for the corresponding view. For each time point we obtain a single fused image (see Fig. 4d) consisting of 4000 × 2000 × 360 voxels. One fused volume was acquired within 40 seconds, providing sufficient temporal resolution to closely follow cell division (see Fig. 4ef). This spatio-temporal resolution and imaging volume is important because many applications in developmental biology require the ability to track cells and follow lineages of the whole animal – something that is very difficult or nearly impossible if cells divide faster than the imaging limits.

**Figure 4:**
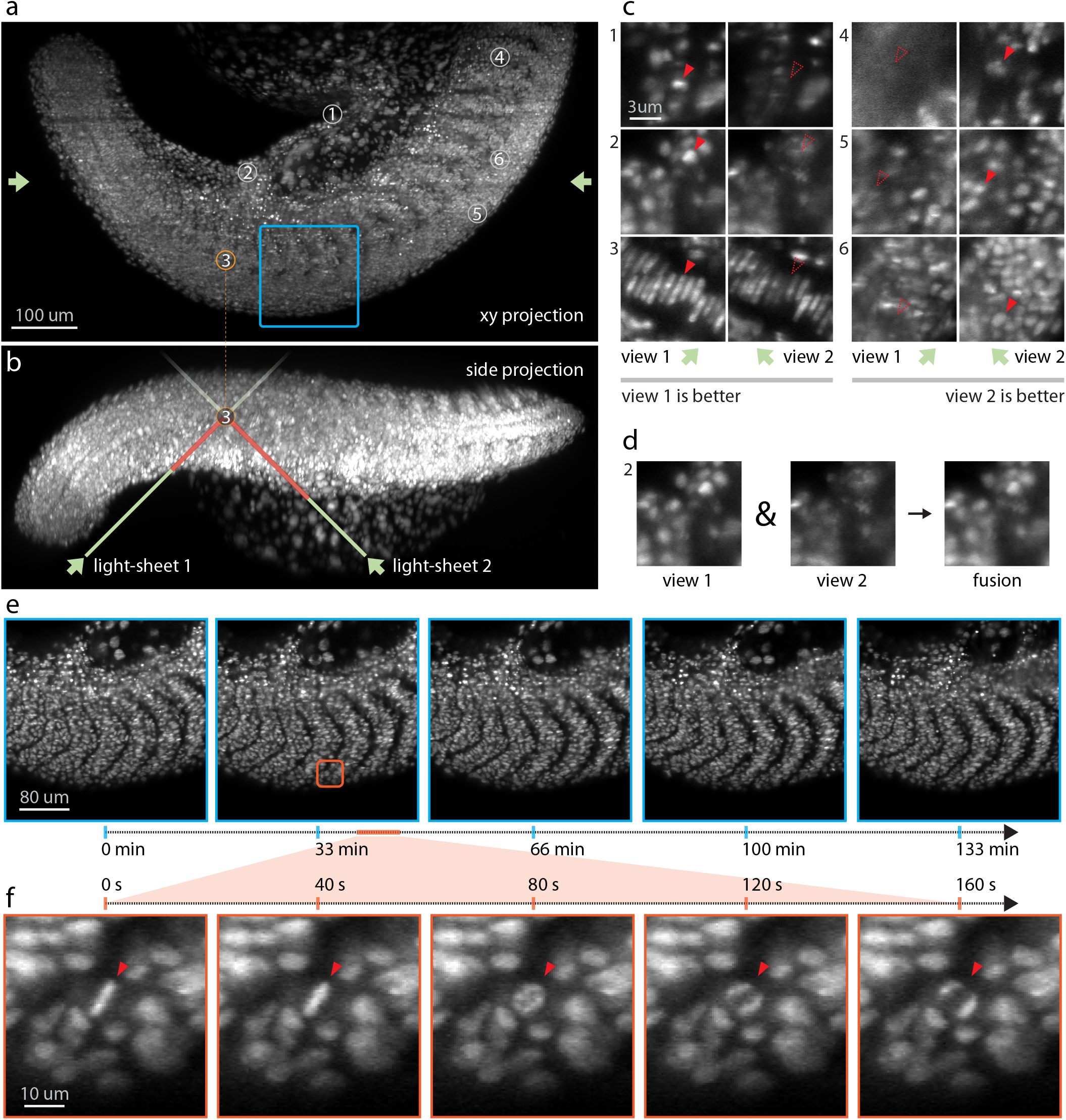
High-speed multi-view imaging of zebrafish tail development. **(a)** Axial max projection showing the whole zebrafish tail at 24 hpf, nuclei labelled with *tg(h2afva:h2afva-mCherry*. Imaging volume is 1064 μm×532 μm×287 μm consisting in 4000 × 2000 × 360 voxels per view for a total of 5.7 billion voxels acquired every 40 seconds. **(b)** Side projection illustrating how the two light-sheets enter the sample at 45° to reach a given point within the sample. Depending on the sample geometry and placement, one of the two light-sheets will have a shorter path to reach that point and hence be less susceptible to absorption, refraction or scattering. Consequently, the corresponding view’s image will be more complete and better contrasted. **(c)** Example regions (single xy plane slices) that demonstrate the complementarity of the two views. In some regions (left) the first view has better image quality, whereas in other regions (right) the second view is better. **(d)** After registration, the two views can be fused together to obtain one high-quality image. **(e)** Time-lapse max-projection frames over a 2.2 hour period centered on the dorsomedial tail, during which time, the boundary between neighbouring somites are accentuated. **(e-f)** Spatio-temporal zoom centered around a cell division, single xy plane slice. Despite the large field of view both views are acquired every 40 seconds making it possible to follow the intermediate steps during mitosis – an important capability for achieving, for example, accurate lineage tracking.

We also recorded somitogenesis during zebrafish development (from 10 hpf to 18hpf), as well as tail extension (from 24hpf to 32hpf) for 8 hours (see Supplementary Fig.11-12 and Movie 3-4). The imaging volume was up to 2200 μm × 800 μm× 300 μm, with each volume captured at 30 second interval. With our microscope, we are able to follow the distinct stages of zebrafish tail development at high spatio-temporal resolution for 8 hours. Longer imaging sessions are conceivable and only limited by the effectiveness of the sample immobilization protocol.

### Simultaneous imaging of nine zebrafish embryos

Lastly, the inverted configuration of the microscope is compatible with multi-well imaging which facilitates simultaneous imaging of multiple samples. Previous attempts at simultaneously imaging multiple samples in a large imaging volume and multi-view light-sheet microscope required the mounting of up to five embryos in FEP tube sections assembled using FEP connectors^21^. While successful, this approach is inherently non-scalable and unpractical as it forces upon the user a delicate sample mounting protocol. In contrast, single-objective light-sheet microscopes let users reuse standard sample mounting protocols already developed for standard microscope systems (e.g. wide-field and confocal). Here, as a proof of concept, we mounted nine embryos in 0.1% agarose gel in a glass-bottom Petri dish (Fig. 5a) and imaged them sequentially (at ~ 4.5 minutes per round) for a total of 8 hours (Fig. 5b-c and Supplementary Movie 5). The samples were mounted in an orientation that allowed us to focus on the embryos posterior development. Overall, the simultaneous imaging of the nine samples was very reliable and provided extremely good resolution, allowing us to follow developmental processes such as somitogenesis in each embryo at single-cell resolution. This mounting strategy is easily scalable, allowing for simultaneous imaging of multiple samples and high throughput screening.

**Figure 5:**
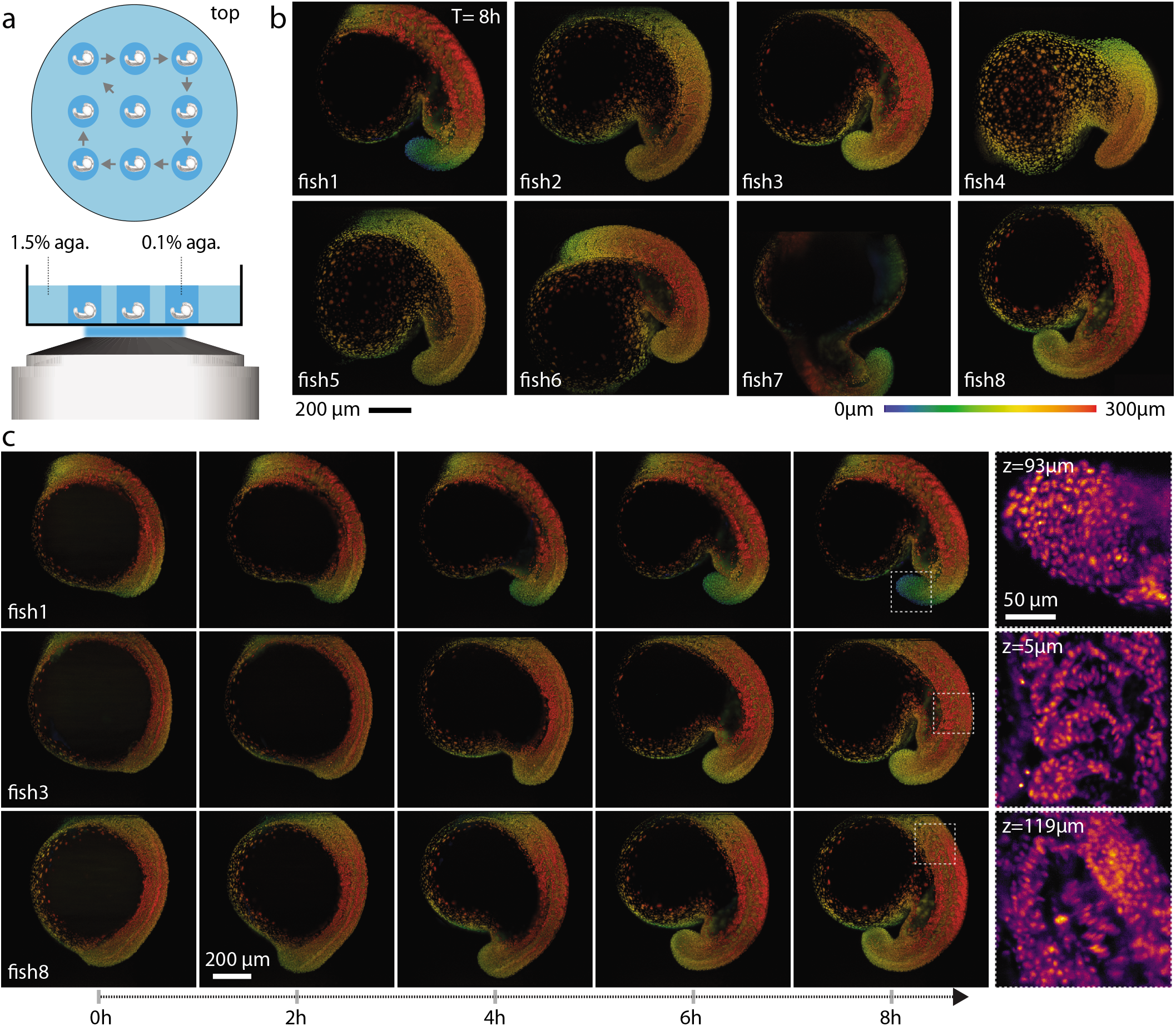
Imaging nine zebrafish embryos at a time. **(a)** Top and side views of nine zebrafish embryos mounted in 0.1% agarose gel. **(b)** The embryos (only eight are shown) were imaged sequentially (at 4.5 minutes/round) for up to 8 hours. Only the final frame at t = 8h imaging are shown (see also Supplementary Movie 5 for the time lapse of all nine embryos). All the embryos developed normally. The images are max intensity projection, color-coded for depth, of the 3D volume. **(c)** Five time points from three different fishes are shown, illustrating the imaging reproducibility across multiple samples.

## 3 Discussions

We have designed, built and characterized DaXi, a novel single objective light-sheet microscope that can achieve high spatiotemporal resolution over a large imaging volume, and long scanning range without introducing motion blur or sacrificing imaging speed. Moreover, our microscope allows for improved illumination coverage through dual illumination/view imaging. We achieved a resolution of ~ 450 nm laterally and 2 *μ*m axially and an imaging volume of 3000*μm* × 800*μm* × 300*μm* (first dimension is only constrained by the stage travel range which is 75 mm with our stage). Previous OPM systems with full NA detection^12,14^ reported an imaging volume of 180*um* × 180*um* × 60*um*, whereas DaXi can image a volume about 400 times larger. Compared to SCAPE^20^ which uses the same primary objective and reported an effective detection NA of 0.35, our microscope achieves full NA detection through refractive-index-mismatched remote focusing and has a detection NA close to that of the primary objective, i.e. 1.0.

The novel light-sheet scanning approach, LS^3^, extends the scanning range of the system by more than a factor of ten, while maintaining high temporal resolution without inducing motion blur. Suppression of scattering and shadowing artefacts is achieved by illuminating and detecting views from two orthogonal directions, following a similar rationale as other multiview light-sheet microscopes^15,16^, but implemented in a single objective geometry. We demonstrated the capabilities of our microscope by imaging large samples, including zebrafish larvae and *Drosophila* fly egg chambers without the need for tiled acquisition. The versatile mounting strategies, low phototoxicity of light-sheet optics and high spatio-temporal resolution of the microscope design allows us to image zebrafish embryo development for 8 hours at 30 second time intervals. In combination, these features will allow investigators to precisely follow cell fate and lineages with improved accuracy, pushing the limit of imaging to enable the capture of complex cellular choreography during early embryonic development.

Crucially, DaXi achieves this performance without forcing unconventional sample mounting procedures on the user – especially when compared with standard multi-view light-sheet microscopes^16,18^. We introduced the remote coverslip method to convert our microscope from upright to a more versatile inverted configuration, which is compatible with many imaging modalities. Biologist can now mount samples in a convenient manner, including multi-well plates, prepare multiple specimens only to image the best, image multiple specimens in parallel, flow drugs and other chemicals with microfluidics, while maintaining a sterile environment. The free space above the sample can potentially be leveraged to combine imaging with other sensing or manipulation schemes such as patch clamping and mechanical perturbation. We demonstrated parallel imaging of nine samples in this paper as a proof of principle. The number of samples that can be imaged/screened can be largely increased using a multi-well mounting protocol^11,12^. Augmenting our system with a microfluidics sample mounting system would enable large-scale screens of mutants or drug-treated animals.

Many of the concepts introduced in this paper are also applicable to other imaging modalities. The remote coverslip method can easily convert objective lenses between immersion and dipping using remote focusing, expanding their utility^22^. LS^3^ can also be used in other light-sheet systems to speed up imaging and to increase the scanning range, by adding a fast actuator such as a galvo to actuate the light-sheet. The dual illumination further improves the overall image quality by providing alternate illumination and viewing angles. Potentially, image rotation by angles different from 180 degrees could add even more illumination and detection diversity. The dual view image flipper is not limited to OPM with full-NA detection, but is also applicable to other variants of OPM^10,23,24^. Sparks et al. independently reported another method for dual-view imaging in OPM, by translating a pair of tilted mirrors in a folded remote-focusing setup^25^. Both methods have their pros and cons, where our image flipper can be integrated without a light loss into an existing system.

The spatial resolution of the microscope can be further improved through adaptive optics^26,27^ to correct both the system and sample-induced aberration. Currently, the two views are fused through content-aware blending by picking from either view for each location of the imaging volume. It is also possible to further improve the resolution through multi-view deconvolution^25,28,29^.

Given its versatility, we expect the microscope described here to be broadly adopted by researchers and applied to a wide range of multi-scale and high-throughput imaging applications, including organoids, which are particularly suited to multi-well imaging and cleared tissue^30,31^ where the high spatio-temporal resolution and large imaging volume would prove to be critical. Future development directions, including smart microscopy and data processing with machine learning, will further improve the optical performance of the system, the versatility and ease of use of the microscope, and the data processing efficiency and quality.

## 4 Methods

### Optical setup and characterization

Supplementary Fig. 1 shows the detailed optical setup. A primary objective (*O*_1_, Olympus XLUMPLFLN 20XW NA1.0, water) is used to both generate an oblique light-sheet in the sample and to collect the fluorescence. A series of tube lenses (*TL*_1_ to *TL*_6_) conjugate the pupils of *O*_1_ and *O*_2_ so that an intermediate image of the sample at *O*_1_ is formed at the secondary objective *O*_2_ (Olympus UPLXAPO20X). The intermediate image has a uniform magnification of 1.33, equal to the refractive index ratio of the medium of *O*_1_ versus that of *O*_2_, making the optical train aberration-free over a reasonable volume^19,32^. A custom tertiary objective *O*_3_ (AMS-AGY v2.0, see Supplementary Fig. 2-4 and Supplementary Note 3) is oriented by 45° with respect to *O*_2_. The fluorescence is filtered by either individual bandpass filters (Chroma ET525/50, ET605/70) or a quad-band filter (Chroma ZET405/488/561/640) and then detected by a scientific complementary metal-oxide semiconductor (sCMOS) camera (Hamamatsu ORCAFlash 4.0). The pixel size of the cameras at the sample space is 146*nm* (*TL*_7_ - Thorlabs AC300A) for PSF calibration, and 265*nm* (*TL*_7_ - Thorlabs TTL165-A) or 439*nm* (*TL*_7_ - Thorlabs TTL100-A) for imaging so as to capture the desired field of view. Objective *O*_3_ is mounted on a piezo stage (PI Fast PIFOC Z-Drive PD72Z1SAQ) so that its focus can be finely adjusted. See also Supplementary Note 1-2 for a more detailed description and alignment procedure, Supplementary Fig. 13 for the 3D rendering of the optical setup of the microscope, and Supplementary Movie 6 for an animation of the setup.

The microscope’s PSF was measured using 100 nm green fluorescence beads. The beads are first embedded in agarose gel (0.5%) and then deposited on a glass coverslip (No. 1.5). The resulting images are then deskewed to the objective-aligned frame of reference (xyz coordinates). Images of single beads are then manually selected and cropped. To account for the tilted PSF with respect to the z-axis, the cropped images are rotated within the xz plane so that the long axis of the PSF is along the z-axis. The PSFs are then fitted with a 1D Gaussian function along all three axes. The values of the FWHMs are averaged from six fluorescence beads in the center of the field of view.

### Light-sheet incident angle adjustment and stripe reduction

A two-axis galvo mirrors (Cambridge 10*mm* 6SD12056) are conjugated with the sample plane so that rotating the two mirrors results in a rotation of the excitation beam at the sample plane. In particular, the incident angle of the light-sheet at the focal space of *O*_1_ can be adjusted by one of the mirrors to 45^°^ with respect to the optical axis. The effective excitation NA is estimated to be 0.08 at this incident angle. The effective excitation NA can be potentially increased, either through reducing the incident angle of the light-sheet or using objectives with higher NA. Increasing excitation NA would allow generating a thinner light-sheet to further improve the axial resolution but at the expense of field-of-view^33–35^. To reduce the stripe due to sample absorption and obstruction of the illumination light, the light-sheet is swept continuous from −6^°^ to 6^°^ within the illumination plane during the acquisition of each frame, using the other mirror of the two-axis galvo.

### Multi-view switching

Two switching galvo mirrors are used to create two views of the object at the sample space. By adjusting the angles of the two galvo mirrors, the light is reflected either by two reflective mirrors or only one. The operating principle is similar to that of a Dove prism. Moreover, the switching module also changes the incident angle of the light-sheet between +45° and −45° since the excitation light also passes through this module.

### Illumination and detection scanning

A galvo mirror (Cambridge Tech, 20mm galvo, 6SD12205) is conjugated to the pupil planes of both *O*_1_ and *O*_2_. Rotating the galvometer actuated mirror scans the oblique light-sheet across the sample (along the x axis), with the incident angle kept at 45°. The Galvo mirror also descans the intermediate image at the focal space of *O*_2_ so that the intermediate image is always projected at the focal plane of *O*_3_. Using the galvometer for 3D scanning allows for faster imaging compared to stage scanning – but at the cost of a limited scan range of approximately 300*um*.

### Motorised sample stage

A motorised stage (ASI MS-2000) is used to position the sample and to perform scanning for volumetric imaging. During acquisition of each frame, stage scanning is combined with galvo descanning to stabilise the imaging plane (coplanar light-sheet and detection planes) relative to the sample. In the absence of relative motion between the sample and the imaging plane, no motion blur occurs. Stabilised lightsheet stage scanning allows for much longer ranges compared to galvometer-based scanning. It is only limited by the range of travel of the stage, in our case 75*mm*. Importantly, illumination and detection planes remain fixed and optimally placed at the center of *O*_1_ and *O*_2_ native axes thus guaranteeing optimal light collection, minimal aberrations, and thus optimal image quality.

### Water dispenser for water immersion lens

For long term imaging, a water dispenser was built to automatically supply immersion water between the primary objective and the sample (Supplementary Fig. 14). The water dispenser consists of a micropump (part of a Leica Water Immersion Micro Dispenser), a microcapillary tip (Eppendorf Microloader) and a 3D-printed objective cap.

### Microscope control software

The data acquisition and display is done by the open-source, freely available software *Micro-Manager 2.0 Gamma*^36^. A custom-developed micromanager script sets up the acquisition order and stores the hyperstack data in TIFF files. A NI DAQ system programmed using a Python module controls the timing of all devices during acquisition. The NI DAQ system consists of one compact chassis (cDAQ-9178), two analog control modules (NI 9263 4-Channel AO) and one digital control module (NI 9401 8-Channel DIO). The Python module uses the NI-DAQmx python API to program the DAQ system. It synchronises the devices, including camera, motorised stage, galvo mirrors and lasers during data acquisition. Alternatively, one can also use Pycro-manager^37^ to perform data acquisition in the Python environment instead of the Java environment of Micro-Manager. The water dispenser is controlled by another Python module to supply water to the primary objective during long term recordings.

### Image processing library – *dexp*

All data processing is done using our open-source Python package *dexp* (Dataset Exploration and Processing). This library performs a number of image processing tasks specific to light-sheet imaging including: equalisation, denoising, dehazing, registration, fusion, stabilization, deskewing, deconvolution. It leverages *napari*^38^ for multi-dimensional visualision and 3D rendering, *CuPy*^39^ for GPU-accelerated computing, *Dask*^40^ for scalable array computing, and *zarr*^41^ as multi-dimensional data storage format.

### Image processing pipeline

The multi-dimensional data are saved by Micro-Manager to local storage of the control PC in TIFF format. The TIFF files are then read out and converted to *zarr* format where typically a 5-10 fold data compression is achieved through lossless compression. The *zarr* datasets are then transferred to a local server equipped with 200 TB storage and 4 NVIDIA Tesla V100 SXM2 32 GB for further processing. The data processing pipeline is shown in Supplementary Fig. 15. Each 3D stack from one of the views is deskewed to coverslip-based coordinates, i.e. the xyz coordinates (see Supplementary Fig. 16). The images are then dehazed to remove large-scale background light caused by scattered illumination and out-of-focus light. The image from the second view is then registered to the first view using an iterative multi-scale warping approach (see Supplementary Fig. 17). Within each iteration: the images are divided into chunks along all three axes by a factor of 2^i^ where *i* is the current iteration number; corresponding chunks from both views are registered separately with a translation model to produce a translation vector; a vector field is then calculated based on all the translation vectors; the image of the second view is warped according to the vector field. This procedure repeats until the max number of iteration or the minimal size of the chunk is reached. In this work, the max iteration number was set to 4 and the minimal chunk size 32 × 32 × 32. This procedure results in a spline-interpolated vector field that is applied to the second view. After registration, images from the two views are fused by picking regions from one or the other image based on the local image quality^42^. The magnitude of the Sobel gradient was used as the metric to generate a blend map, based on which the two images are blended. Other fusion methods such as frequency domain fusion (DCT, FFT) are available in *dexp*. An illumination intensity correction is applied to the fused data to account for the Gaussian profile of the light-sheet along the y-axis. Each raw data per time point is ~ 8GB (two views, ~ 1000*1024*2048*2 voxels). An 8 hour continuous imaging session produces ~ 8 TB raw data (~ 1000 timepoints). It takes ~ 1 minutes to process the data per time point per GPU, 4 hours to process the whole dataset with 4 GPU running in parallel. After all time points are processed, they are temporally stabilised to compensate minor drifts overtime.

### Sample preparation

Zebrafish husbandry and experiments were conducted according to protocols approved by the UCSF Institutional Animal Care Use Committee. In the experiments we used the *tg(h2afva:h2afva-mCherry* (a gift from Jan Huisken, Morgridge Institute for Research, Madison, USA) and for the functional imaging of neuronal activity the *tg(elevl3:GCaM6f)* and the *tg(elavl3:H2B-GCaMP6s)^43^*. The sample mounting geometry is shown in Supplementary Fig. 18. First, zebrafish were dechorionated with a pair of sharp forceps underneath a binocular dissecting microscope, and incubated at least 5 minutes in a solution of fish water and tricaine (0.016%). Embryos were gently pipetted into a 0.1% solution of low gelling point agarose (Sigma, A0701) cooled at 37°*C*. The embryos, together with approx. 1*ml* of 0.1% agarose, were placed in a glass-bottomed petri dish (Stellar Scientific Cat. No 801001). Using a capillary needle, the embryo is gently positioned at the center of the dish and in the desired orientation, here laterally. When the agarose is solidified, the dish is flooded with fish water and 0.016% tricaine. The sample mounting geometry is also shown in Supplementary Fig. 19. All the time lapse experiment were done with a gentle flow of embryo medium water with 0.016% tricaine at 28C, using peri-staltic pumps, allowing normal development of the embryos. For the calcium imaging experiment fish were embedded in 2% agarose and anaesthetised in external solution with tricaine (0.2 mg/mL) at 5 dpf. For Drosophila fly imaging, the egg chambers were dissected and cultured as described previously^44^. The egg chambers are then transferred to a glass-bottomed petri dish for imaging. The mounting is similar to the zebrafish imaging, except that the egg chambers are immersed in imaging media rather than agarose solution.

## Data availability

All datasets acquired will be made available upon request and will be eventually deposited in a public repository.

## Code availability

All code can be found at github.com/royerlab/daxi. These include the python code for optical simulation, device control and the Micro-Manager acquisition script. Our image processing tool **dexp** (Dataset Exploration and Processing Tool) can be found at github.com/royerlab/dexp.

## Supplementary Information

All supplementary information, code, blueprints and tutorials can be found at: *github.com/royerlab/hr-lf-mv-SOLS*

## Author Contributions

BY and LAR conceived the research. BY designed and built the microscope, and performed all imaging. BY wrote the control software with help from ACS. AMS and AGY designed the Snouty 2.0 objective. BY, AGY and RPF designed the image flipper for dual view imaging. LAR and BY wrote the image processing code. ML designed the sample mounting protocol and prepared the zebrafish samples with help from SV. WW prepared the Drosphila fly samples with guidance from TBK. MNM provided the zebrafish samples for brain imaging. LWW made the visual abstract video. BY, ML, HK, and LAR analysed the data. BY and LAR wrote the paper with input from all authors.

## Acknowledgements

We would like to thank the Bioengineering team at CZ Biohub for loaning us the water immersion micro dispenser. We thank Nico Sturrman from UCSF for his help on Micro-Manager and Jan Huisken (Morgridge Institute for Research) for sharing the Tg(h2afva:h2afva-mch) line. Sincere thanks go to Ashley Lakoduk, Mirella Bucci, and Sandra Schmid for their constructive criticism and diligent proofreading of the manuscript. Funding for this work was provided by the Chan Zuckerberg Biohub. We thank Calico Life Sciences LLC for funding the development of the AMS-AGY V2.0 objective and for granting a loaned objective.

## Competing Interests

A patent application has been filed covering the reported microscope design.

## Notes

https://github.com/royerlab/daxi

### Summary of Updates

This is an updated version of the manuscript that reports on more imaging experiments.

