## Supplementary Information for "High-Resolution, Large Imaging Volume, and Multi-View Single Objective Light-Sheet Microscopy"

### 1 Supplementary Note – Optical setup

Supp. Fig. 1a shows a detailed scheme of the optical setup of the microscope. A primary objective (O1, Olympus XLUMPLFLN 20XW NA1.0, water) was used to both generate an oblique light sheet in the sample and to collect the fluorescence. A series of tube lenses (TL1 - TL6) conjugate the pupil pupils of O1 and O2 so that an intermediate image of the sample at the focal space of O1 was formed at the focal space of the secondary O2 (Olympus UPLXAPO20X). The intermediate image has a uniform magnification of 1.33, equalling to the refractive index ratio of that of O1 and O2, so that it is aberration-free. A tertiary objective O3 (Calico AMS 2.0) is oriented by 45° with respect to O2. The fluorescence was filtered by either individual bandpass filters (Chroma ET525/50, ET605/70) or a quad-band filter (Chroma ZET405/488/561/640) and then detected by a scientific complementary metal-oxide semiconductor (sCMOS) camera (Hamamatsu ORCAFlash 4.0). The pixel size of the cameras at the sample space was 219 nm (TL7 - Thorlabs TTL200-A) for PSF calibration and 266 nm (TL7 - Thorlabs TTL165-A) for imaging to be able to capture the desired field of view. O3 was mounted on a piezo stage (PI Fast PIFOC Z-Drive PD72Z1SAQ) so that its focus could be finely tuned.

**Illumination.** The illumination light came out of a fiber output which input is from a custom laser combiner of two lasers (Vortran Stradus 488 nm and 561 nm). The light is firstly collimated by a telescope composed of two achromatic lenses (L1-L2) and then expanded along the horizontal direction by two cylindrical lenses (CL1-CL2). It is further focused on to the 2-axes galvo mirrors by CL3 and then reflected by a dichroic mirror (Chroma ZT405/488/561/640rpc) to be combined with the detection path. The 2-axes galvo mirrors (Cambridge 10 mm 6SD12056) are conjugated with the sample plane so that rotating the two mirrors resulted in a rotation of the excitation beam at the sample plane. In particular, the incident angle of the light sheet at the focal space of O1 can be adjusted by one of the mirrors to 45° with respect to the optical axis. The effective excitation NA is estimated to be about 0.08.

**Optical reflector.** The two switching galvo mirrors shown in Supp. Fig. 1b can create two views at the remote space of the object at the focal space of O1. By adjusting the angles of the two galvo mirrors, the light is reflected either by M5 and M7 or by M6 only. The operating principle is similar to that of a Dove prism. The red arrow is reflected one more time compared to the green arrow. As a result, it is flipped along one direction compared to the green one. The intermediate image of the sample is therefore flipped along the horizontal plane. Supp. Fig. 1c shows two images of a calibration grid taken with the two views, and it clearly shows that the image in the right view is flipped along the horizontal axis compared to the image shown on the left. Moreover, the switching module also changes the incident angle of the light sheet between +45° and -45° since the excitation light also passes through this module. Instead of using gavlo mirrors, one can also consider using a mirror mounted on a motorised stage to send the light to different paths. We chose galvos mirrors for these purposes since they are often one order of magnitude or even more faster than motorised stage. It is also possible to have a mirror fold system mounted on a rotating mount. By rotating this module, one would be able to create more than two views of the sample, with in principle unlimited possible views.

**Imaging plane scanning.** A Galvo mirror (Cambridge Tech, 20mm galvo, 6SD12205) was conjugated to the pupil planes of both O1 and O2. Rotating the Galvo mirror scanned the oblique light sheet across the sample (along the x axis), with the incident angle kept at 45°. The Galvo mirror also descanned the intermediate image at the focal space of O2 so that the intermediate image was always projected at the focal plane of O3. Using the galvo for image scan allows faster imaging speed compared to stage scanning. The scanning range is limited to

about 300 μm, both due to cropping of the excitation beam by the relay tube lenses and decreased optical performance when the illuminated plane is away from the optical axis of O1.

### 2 Supplementary Note – Optical system construction and alignment procedure

#### Test samples

1. A multi-frequency grid distortion target (Thorlabs, R1L1S1P) to measure the magnification of the system;
2. A #1.5 glass coverslip (170 um thickness) uniformly coated with the fluorescent dye to characterise the light sheet;
3. 170 nm fluorescent beads embedded in 2% agarose gel to characterise the resolution of the system.

#### General remarks and guidelines for alignment

1. There are a few planes are conjugated with 4f-systems in the optical setup. Firstly, the pupil planes of O1, O2, the scanning galvo mirror and the slit are conjugated. The conjugation between the pupil planes of O1 and O2 ensures aberration-free imaging; the conjugation between the scanning galvo mirror and the pupil plane of O1 ensures constant incident angle of the oblique light sheet and proper image descanning; the conjugation between the pupil plane of O1 and the slit allows adjusting the light sheet thickness. Secondly, the 2-axes galvo mirrors and the focal plane of O1 are conjugated so that the incident angles of the light sheet can be adjusted.
2. O1, O2 and O3 are all mounted on translation stages (Thorlabs XR25P) so that their position can be precisely adjusted.
3. Start by placing and aligning all the mirros at their corresponding locations (either through estimation or follow a CAD design).
4. Then place the lenses one by one, using a reflective mirror at the sample plane to adjust the lateral positions, check the light collimation (with a shear interferometer (Thorlabs SI050) or other means) to adjust the axial positions.
5. Lastly fine tune the system to have diffraction-limited imaging, mostly by assuring proper conjugation between critical planes.

#### Detailed alignment procedure

##### Set up all optics

1. Setup the mount for the fibre collimator at desired height.
2. Place mirrors (including the dichroic mirror) after the collimator one by one, ensures that the height is constant and the beam hits the centre of the mirrors. Note that the height is different before and after the two-axes galvo mirrors. Ignore M6 for now, but make sure that the switching galvos

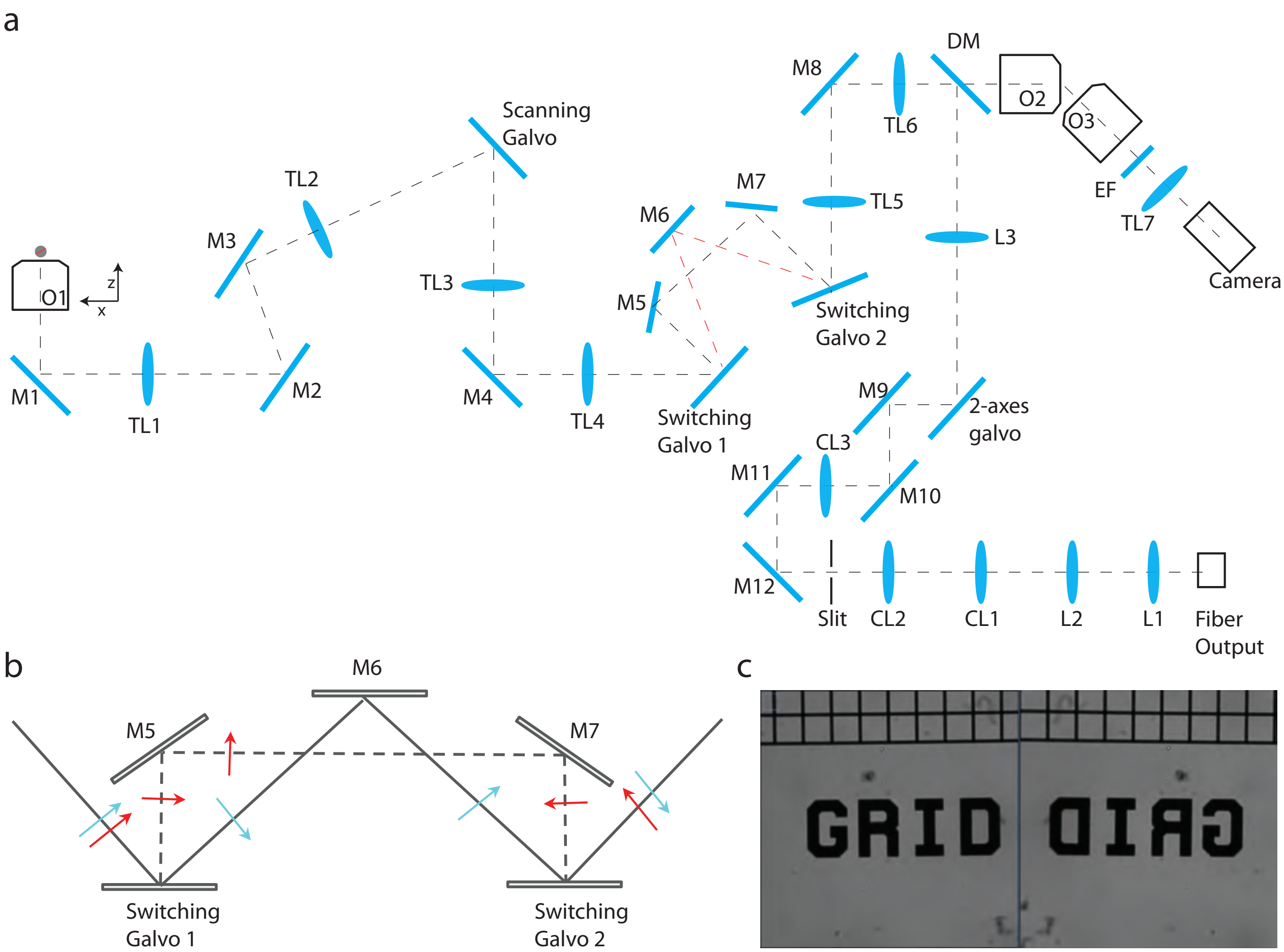


Figure 1: Optical setup of the microscope. **(a)** Detailed layout of the setup. Objectives lenses: O1 - Olympus XLUMPLFLN 20XW, O2 - Olympus UPLXAPO20X, O3 - Calico AMS-AGY v2.0. Tube lenses: TL1, TL2 and TL3 - Olympus SWTLU-C 180 mm, TL4, TL5 and TL6 - 135 mm custom designed tube lens (using off-the-shelf pieces from Thorlabs, see [ref]), TL7- Thorlabs TTL200-A or TTL165-A. Scanning galvo and switching galvos: Cambridge 6SD12205 20mm galvo mirrors. Cylindrical lenses: CL1-CL3. Achromatic double lenses: L1-L2. Dichroich mirror: DM - Chroma ZT405/488/561/640rpcv2-UF3. Mirrors: M1-M12, protected-silver coated mirror. 2-axes galvo: Cambridge 6SD12056 10 mm galvo mirrors. EF: emission filters, Chroma ET525/50m or ET605/75m. **(b)** Dual view switching module. The two switching galvo mirrors can switch the light path so that light is reflected either by M5 and M7 or by M6 only. The red arrow is reflected four times along the path and maintains upright, while the green arrow is reflected only three times and becomes inverted. **(c)** Dual view images of a calibration grid (Thorlabs R1L1S1P). The two images are clearly flipped with respect to each other. The images were taken under bright field illumination and with O3, TL7 and the camera on a straight line with O2.

are set to the correct value and their value can be adjusted to send light to approximately the centre between M5 and M7.

3. Place a few irises (Thorlabs ID25 and ID50) along the path to aid alignment of lateral positions of the lenses.
4. Make sure that the light passes through the mount of O1 centered.
5. Place a reflective mirror at the place of O1, slightly adjust M1 so that the reflected beam passes all the iris centred. Go back to step 4 if necessary.
6. Place L1 and L2 to have a collimated beam.
7. Place TL1, adjust its lateral position so that the reflected beam passes all the iris and its axial location is approximately at the desired location.
8. Place TL2, adjust its lateral position so that the reflected beam passes all the iris, adjust its axial position so that the beam is collimated after TL1.
9. Repeat step 8 for TL3 to TL6.
10. Start with having O2 and O3 along a straight line. Make sure the reflected light passes through the centre of the mounts of O2 an O3, then place O2 and O3 in the setup.
11. Place TL7 in the setup, make sure that the light passes through its centre.
12. Place the camera in the setup, depending on the specificity of TL7, make sure the distance from TL7 to the camera sensor is correct.
13. Place O1 in the setup.

**Adjust the position of O1 and O2 to have them both conjugated to the galvo mirror.**

14. Send a sinus wave signal to the scanning galvo, adjust axially O1 so that the light coming out of it is scanning laterally without changing angles. This is the desired position for O1.
15. Put the grid sample on the sample stage. If necessary, add a spacer beneath it so that O1 is approximately at its desired position.
16. Use a flash light to illuminate the grid and have its image on the camera.
17. Move O3 200 $\mu m$ away and closer to O2, and measure the magnification of the system with the grid. The goal is to have the same magnification regardless of the relative position of O3 to O2. Move O2 and O3 together axially until at one point the magnification is uniform. The pupil plane of O2 is then conjugate to that of O1.
18. Translate the stages to move O2 and O3 together axially until at one point the magnification is uniform.
19. Place a sample with beads imbedded in 0.5% agarose on the sample stage. Place the emission filter, switch on the laser and observe the fluorescence image of the beads. The defocused image of the beads should look circular, otherwise slightly adjust O2 laterally to have a circular image of the beads. Adjust O3 slight to bring back the field of view to the centre of the camera. One can move O3 axially to inspect the images of the beads along different z. The image quality should be constant across at least 500 $\mu m$, otherwise the alignment is subject to further trouble shooting and improvement.

**Dual view component alignment**.

20. Adjust the voltages sent to the switching mirrors so that the light is reflect to the centre of the M6 and also passes through the irises from M6 to O1 thought the centre.
21. Place the grid back to the sample stage and measure the magnification with O3 at 200 $\mu m$ away and closer to O2. Translate the stage to move M6 so that the magnification is uniform when translating O3 axially.
22. Repeat step 20.

**Oblique light sheet setup and alignment**.

23. Place O3 and the components downstream at 45° to O2. Place a sample with beads imbedded in 0.5% agarose on the sample stage. Adjust O3 so that the fluorescence image of the beads is capture by the camera in the centre of the field of view.
24. Set up CL1 and CL2 so that the laser light is expanded along the direction horizontal to the optical table.
25. Set up CL3 so that the beam is focused on the 2-axes galvo.
26. Place the slit at approximately the focal plane of CL3.
27. Place L3 so that the light becomes collimated again along the vertical direction.
28. Place a sample with beads imbedded in 0.5% agarose on the sample stage. Adjust the sample of the y-axes galvo to adjust the incident angle of the light sheet to 45° within the xy plane. Adjust O3 slightly to refocus if necessary. When the light sheet is at the correct angle, all beads in the image will appear in focus.

### 3 Supplementary Note – AMS-AGY v2.0 Objective – Full technical details and context

Light-sheet fluorescence microscopy (LSFM) or selective plane illumination microscopy (SPIM) is a powerful technique for biological imaging. Fast, gentle, and with good sectioning, the idea has seen much attention with innovations like the DiSPIM and Lattice, and commercial instruments like the Zeiss Lightsheet 7 and the Leica DLS. However, traditional light-sheet designs require two or more orthogonal lenses for illumination and collection, resulting in an awkward interface between the biology and the optics, and a major drawback for many users and applications.

The Oblique Plane Microscope (OPM) invention of 2008 restored the traditional coverslip boundary by passing light-sheet excitation and emission through a single primary objective, and then using a tilted remote refocus (RR) in the downstream optics to image the equally tilted plane of illumination (the object plane). OPM showed that light-sheet microscopy could be done with a standard microscope and sample interface, but seemingly exchanged this convenience for heavy losses in resolution and optical efficiency.

The problem with OPM was that the tilted portion of the remote refocus would lose a significant fraction of the emission light, simply because the numerical aperture of the final objective was too low. An ideal objective for this location would collect all of the emission light, even with the additional tilt imposed by the OPM architecture, i.e. a full hemisphere of collection (a seemingly impossible requirement when objective half angles are typically limited to 70 deg).

However, in 2018 the Epi-illumination SPIM microscope (eSPIM) showed that the major optical losses in OPM-style systems could in fact be avoided. By using a water objective and coverslip assembly as the final objective, the numerical aperture could now equal the refractive index of air (1.0) i.e. the 'immersion medium' of the opposing objective in the remote refocus. This is the crucial insight: to make the elusive 'hemisphere' collection objective you simply need a numerical aperture that is greater than (or equal to) the index of the medium in which it operates. So for example, an NA 1.0 water immersion objective has a modest 49 deg collection cone in water (a reasonable lens to manufacture), but in air this transforms to 90 deg. So by imaging at the coverslip boundary, a water lens can indeed collect a solid angle of 2pi from an air medium.

There are however some additional considerations that complicate the eSPIM approach. The tertiary objective assembly is corrected for a water/coverslip/water medium (not water/coverslip/air). It can perform well if operated exactly at the surface of the coverslip, but deviations in alignment that push the image into the coverslip (or out into the air) will produce strong aberrations, so it can be challenging to align and keep stable (and hydrated). In addition, the bulkiness of the coverslip-water assembly requires longer working distance objectives in the remote refocus to avoid mechanical collision from the tilt. In practice this limits the choice of optics (and tilt range) and can force a reduced numerical aperture on the air objective.

Inspired by OPM and eSPIM, the AMS-AGY v1.0 objective (aka Snouty) was developed to eliminate the previous trade-offs and compress the opto-mechanical difficulty into a single dedicated component (Fig. 2B). The NA 1.0 objective features a monolithic glass tip with zero working distance; this tip is optically equivalent to an oil/coverslip/air interface but alignment-free and mechanically stable. The tip also features an anti-reflection coating to maximize collection from the full hemisphere of rays (as noted previously NA 1.0 in an air collects from the full 90 deg half-angle). The zero working distance is another critical feature; the large refractive index mismatch at the glass-air interface produces strong spherical aberrations that vanish only at this boundary. The high refractive index of the glass tip compresses the collection half-angle so the tip can be shaved off (Fig. 3B) to allow a range of tilt angles from 0-45deg. This excellent mechanical clearance allows image collection as close as 100um from a planar boundary. In practice this means the AMS-AGY objective can be paired with objectives with the highest numerical apertures and therefore the maximum collection efficiency. Infinity and color corrected, this component enables an extensive suite of design options as detailed in the High NA single-objective light-sheet (SOLS) article of 2019.

The Snouty v1.0 objective enabled 'bolt-on' SOLS designs with uncompromised numerical aperture, and is the ideal microscope for many light-sheet applications. However, the design of the v1.0 lens was constrained by mechanical and economic considerations, ultimately limiting the field of view (FOV) to 150um diffraction-limited (250um to the shaved edge, Fig.3B). The economic argument is obvious: manufacturing difficulty and cost typically increase with field of view, and a costly lens would increase the prototyping risk and lower uptake of the technology. The mechanical limitations are more subtle; for the highest numerical aperture SOLS designs, the air objective that opposes the Snouty lens in the remote refocus can have working distances as short as 200um (Fig. 3B). So as the Snouty lens is tilted, the field of view (housed in glass) moves towards a collision with the opposing lens i.e. the Snouty FOV competes directly with the working distance of the paired objective in the remote refocus.

To overcome the limits on field of view the Snouty v2.0 prototype (aka KingSnout) was developed with a 3x boost on FOV compared to the v1.0 lens i.e. 450um diffraction limited and 450um to the shaved edge (Fig 1A, 2A). The large field was partly achieved by increasing the budget for design and manufacture (it is a larger and more complex lens) but also by eliminating the margin between the diffraction limited FOV and the shaved edge. This enables the lens to be used ‘off-axis’ in the tightest of spaces without losing optical quality. Snouty v2.0 is also ground at a more aggressive angle (55 deg vs 45 deg) which combined with the reduced margins gives the maximum clearance for tilting in the remote refocus. It is a strict upgrade to the v1.0 objective and compatible with all SOLS designs.

The discussion of lens specifications brings to the forefront some important considerations that should be emphasised. The AMS-AGY objectives are exactly as specified: NA 1.0 with fields of 150um (v1.0) and 450um (v2.0), where NA 1.0 actually means the lens will image stigmattically at NA 1.0, not that it merely collects at this NA. How these specifications translate into the object space and the resulting volumetric imaging performance is subtle and beyond the scope of this section. However it should be noted that these objectives can be used beyond the specified fields, by up to a factor of approximately 1.7x, but not at NA 1.0. So a less genuine, but perhaps more typical specification would be NA 1.0 with fields of 250um (v1.0) and 750um (v2.0) and is something to bear in mind when considering a SOLS design.

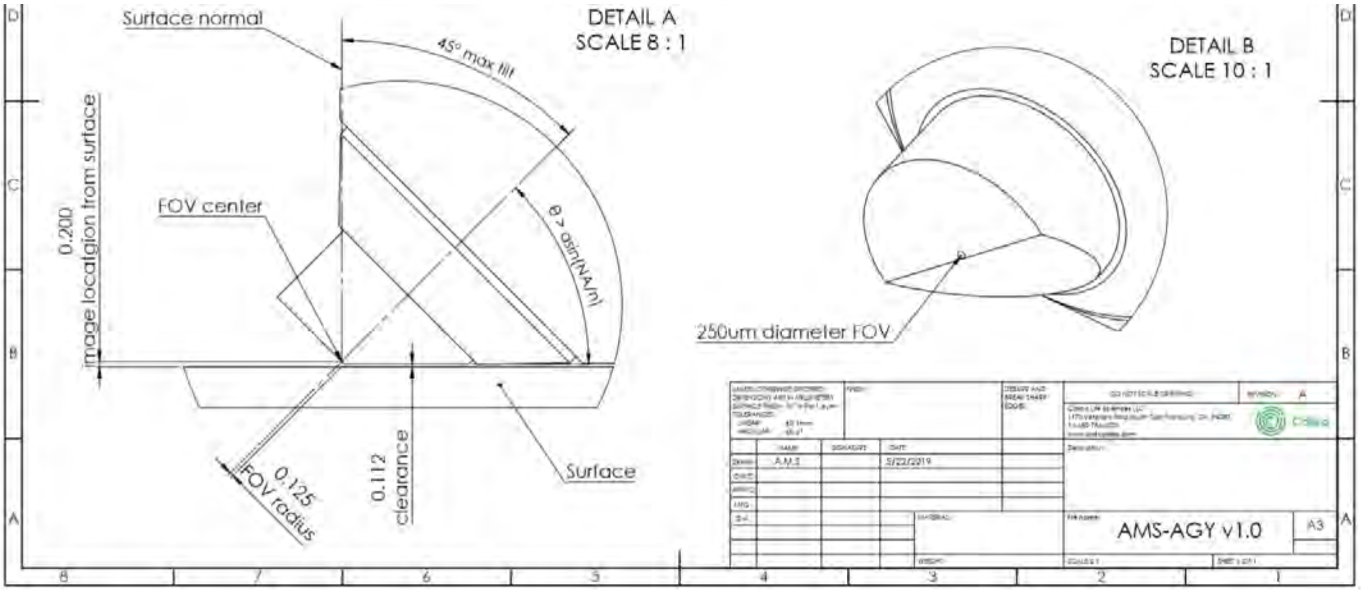


Figure 2: Technical drawing for AMS-AGY V1.0 objective.

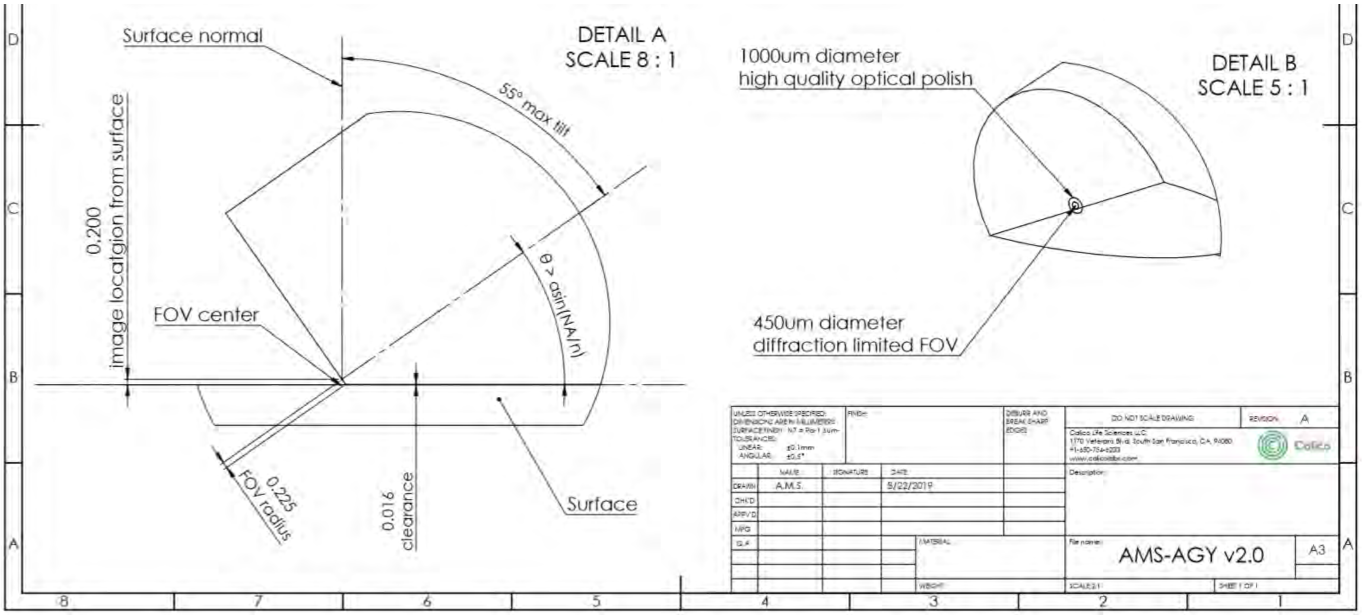


Figure 3: Technical drawing for AMS-AGY V2.0 objective.

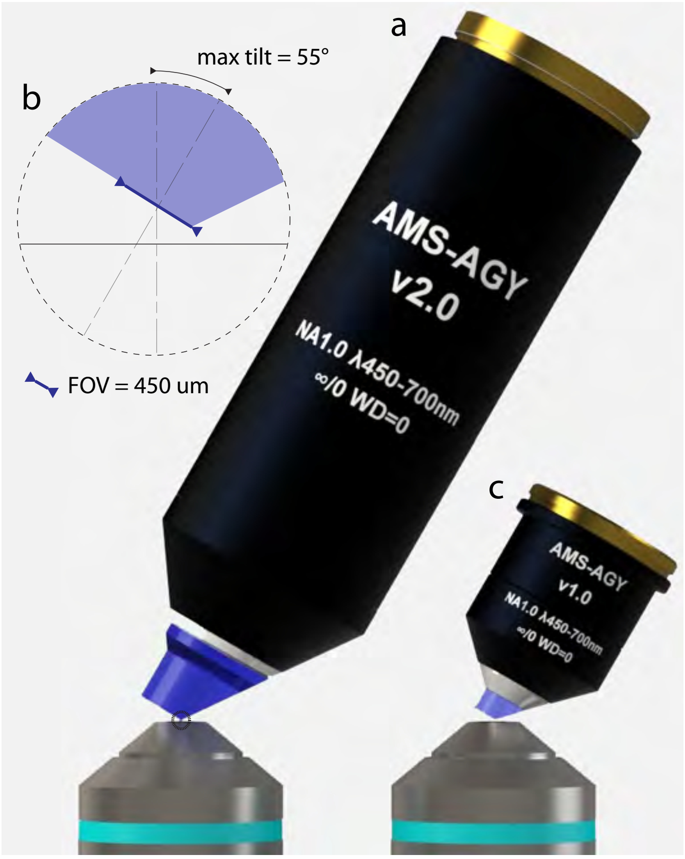


Figure 4: Comparison of the AMS-AGY v1.0 and AMS-AGY v2.0 objectives.

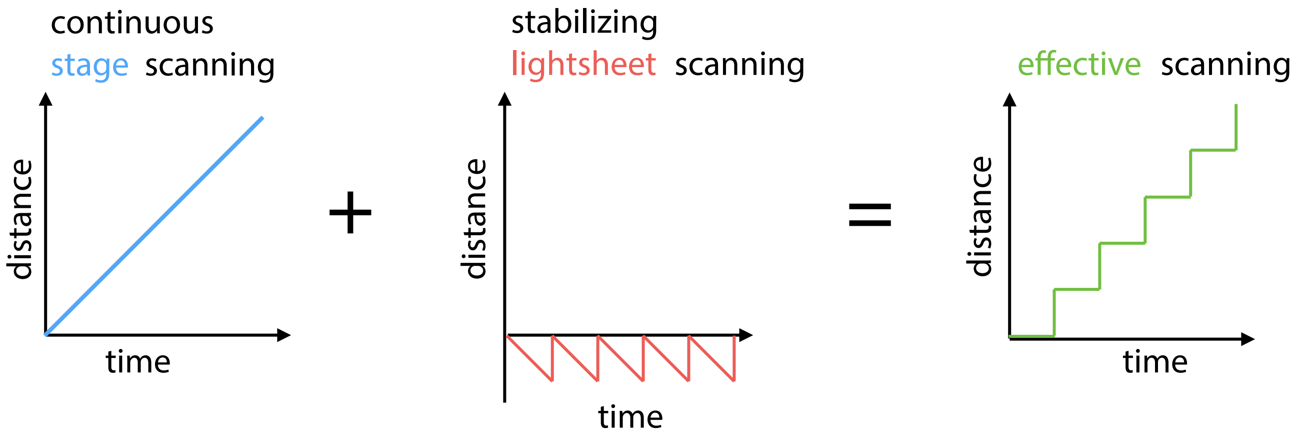


Figure 5: Light sheet stablized stage scanning. During the acquistion of a 3D image stack, the stage moves continuously and the galvometer scanner performs a counteracting motion of the light-sheet and detection planes to cancel out any relative motion between sample and imaging plane, resulting in an effective step-wise scan of the sample.

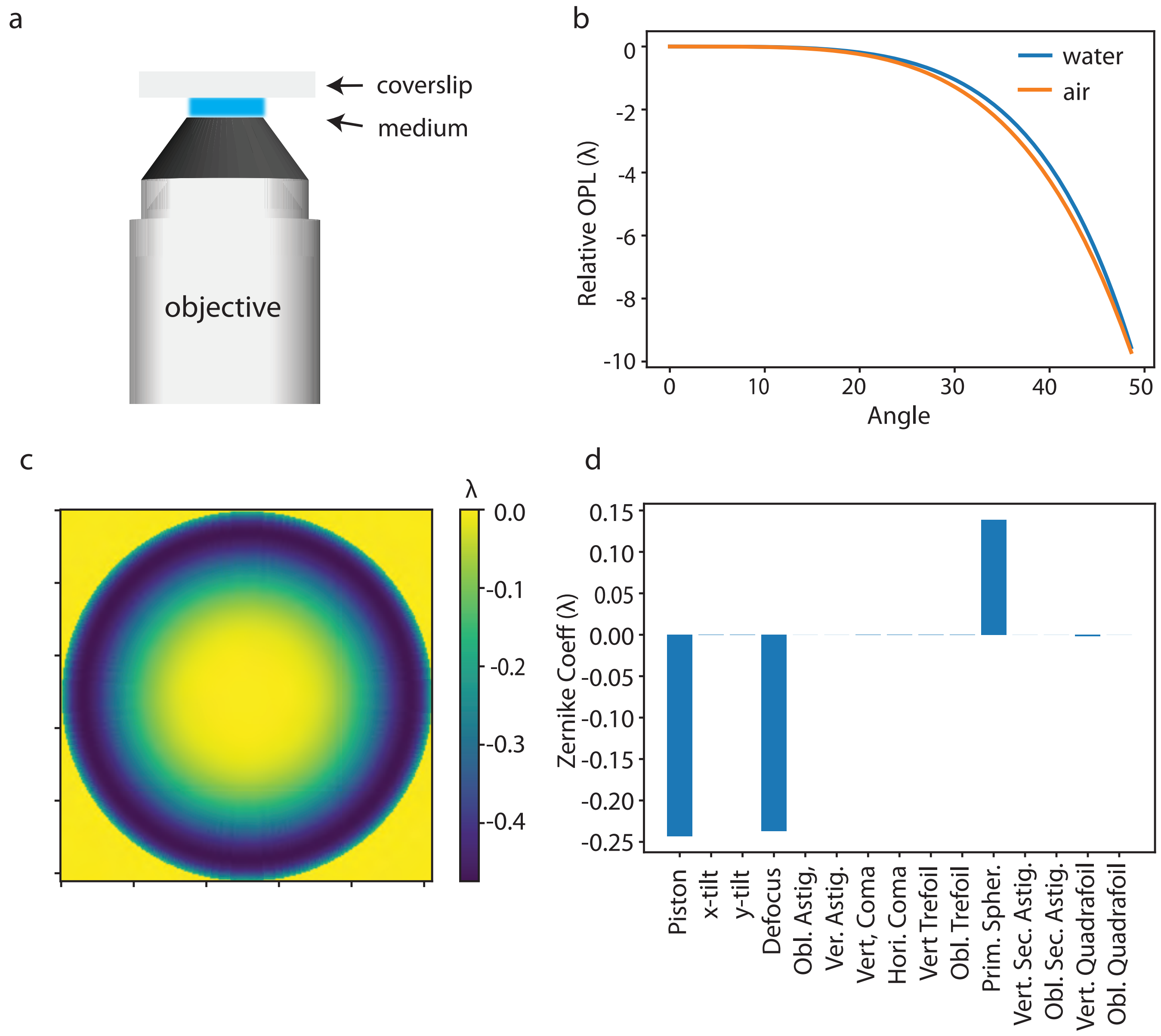


Figure 6: Analysis of the optical path length when placing a glass coverslip into the imaging medium of an objective. **(a)** illustrates the optical system to analyse. Depending on the type of the objective, the medium could be air, water, immersion oil, and etc. **(b)** plots the optical path length of an emitter at the coverslip surface as a function of the angle between the emission ray and the optical axis, in the case of air or water medium. **(c)** shows that 2D map of the optical path difference of using air and water medium. This 2D phase map is then fitted with the first 50 Zernike terms. **(d)** shows the coefficients of the first 20 Zernike terms. The major non-zero Zernike terms are pistol, defocus and primary spherical aberration. This suggests that the primary spherical aberration is the only aberration introduced when the coverslip is moved from the secondary objecrive (air) to the primary objective (water) in a remote focus system (see Fig. 1). Considering that the primary spherical aberration can be compensated by moving any of the relay tube lens in practice, one should be able to maintain the optical performance of the microscope when switching the coverslip's location. Simulation is performed for objectives of NA 1.0 water and NA 0.75 air. The Zernike fitting is done using pyOTF, a simulation software package for modelling optical transfer functions (OTF)/point spread functions (PSF) of optical microscopes written in python. Link: https://github.com/david-hoffman/pyOTF

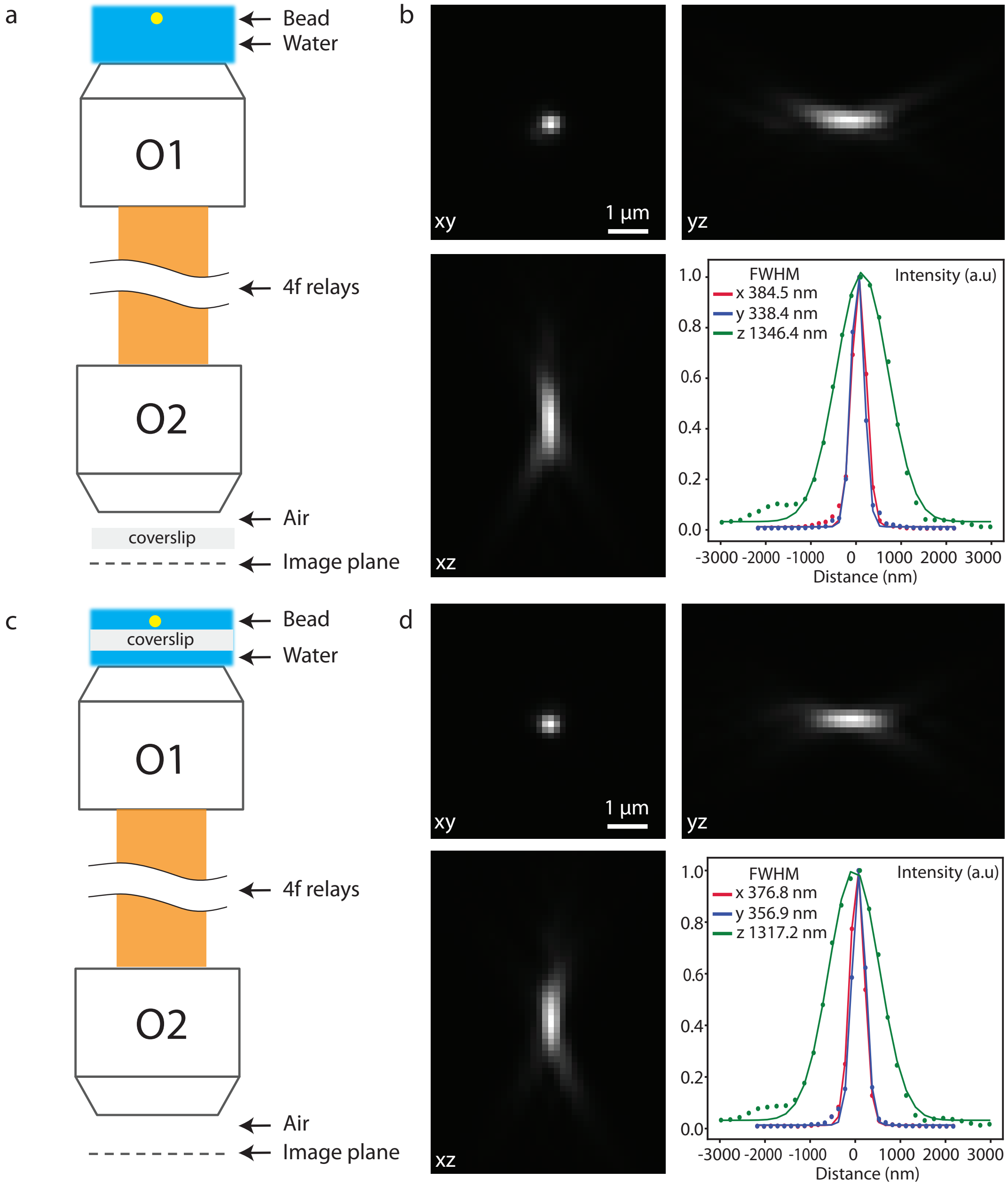


Figure 7: Converting the microscope from upright to inverted by repositioning the coverslip in a remote focusing system. **(a)** shows the remote focusing system composed of two objectives whose pupil planes are conjugated by a 4f relay system. Yellow dot: fluorescent bead. O1: 20x, 1.0NA, water dipping. O2: 20x, 0.8NA, air. According to their factory design, O1 should be used directly facing the sample without any coverslip in between. O2 requires a coverslip between the objective front lens and the image plane, as shown in **(a)**. This configuration is well suited for an upright microscope where the primary objective is used in a dipping configuration. **(b)** shows the PSF of the setup in **(a)** measured with 100 nm green fluorescence beads. Interestingly. According to our analysis (see Supp. Fig. 6), one can reposition the coverslip at the focal space of O2 to that of O1 as shown in **(c)** and can still achieve similar optical performance, turning the system to an inverted configuration. **(d)** shows the PSF of such a system (**c**), which has indeed similar FWHMs along all three axes compared to the values in **(b)**.

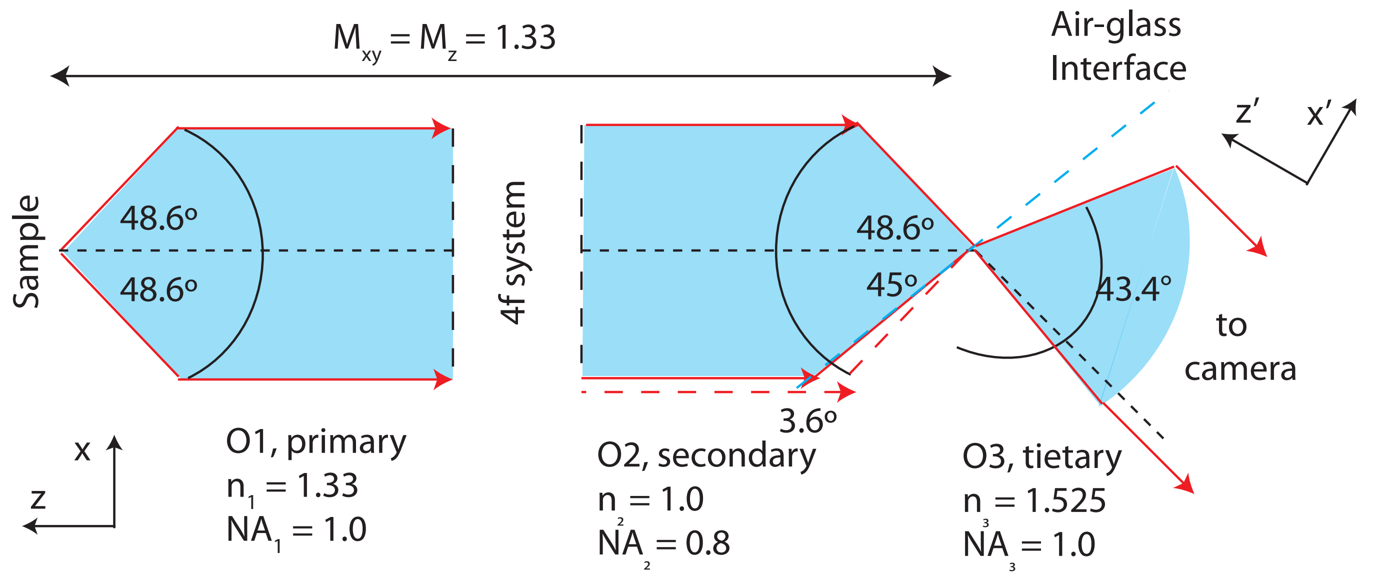


Figure 8: Diagram illustrating the effective detection NA along the x' axis. Blue shades indicate the light cone collected by each objective; red lines represent the light rays with the highest incident angle; black dashed lines show the optical axis. The intermediate image at the focal space of O2 is magnified by 1.33 (equal to $NA_1/NA_2$) along both the lateral and axial directions to minimize the aberration of the intermediate image. The blue dashed line indicates the position of the air–glass interface. All the light passing though O2 is refracted at this interface and enters O3, with the exception of only a small portion of the light being cropped by the interface (left side between the blue and red dashed lines). The effective detection NA along the x'-axis is $1.33 \times 1.525 \times \sin(43.4°/2) \approx 0.75$. The effective NA along the y axis is 1.0, since light is not cropped along this direction.

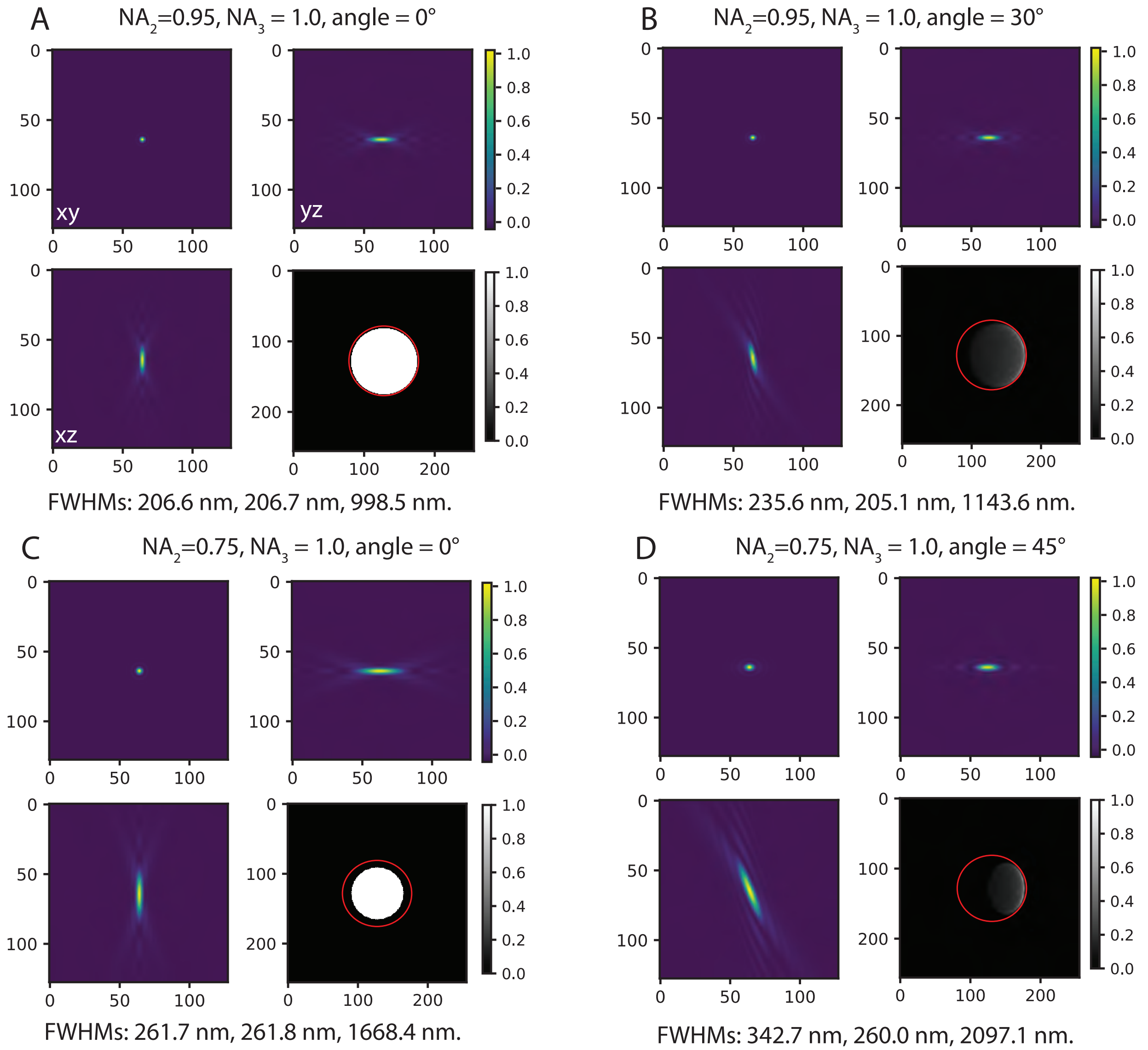


Figure 9: Simulated PSF and pupil function of the eSPIM system. The eSPIM system is simplified to only consider the secondary (O2) and tertiary (O3) objectives. A plane wave is assume to pass through O2 and the pupil function is estimated after the light exits O3. The 3D PSF is then simulated from the pupil function. O2 (NA 0.75 or 0.9) and O3 (NA 1.0) are respectively air and solid immersion objectives. The two objectives are either along a straight line (i.e. angle = 0) or orientated by an angle (30° or 45°) between their optical axes. **(a)** and **(b)** show the simulated PSF (xy, xz and yz cross sections are shown) and pupil function (gray images) for a high NA version of eSPIM, similar to that reported in [1]. The red circles indicate the extend of the pupil of O3. **(c)** and **(d)** show the simulated PSF and pupil function for the optical system presented in this paper. When the angle between the two objectives is not 0°, the effective pupil function of the imaging system show a compression of the light towards one dimension. The resulting PSF is therefore not straight along the z axis, but has an angle to it, which is 10° for **(b)** and 20° for **(d)**. The PSFs are fitted by a 1D Gaussian Function along the three principle axes to obtain the FWHMs, given at the bottom of the images. The tilted PSFs (**b** and **d**) are first rotated to have their principle axes along the x-, y- and z-axes and then fitted with 1D Gaussian Function to get the FWHMs. The FWHMs along y are comparable with O2 and O3 either along a straight line or tilted, suggesting that the effective NA is the same as the secondary objective. The FWHMs along the two other directions are slightly wider when the two objectives are tilted, as also suggested by the analysis in Fig.8. Note that the FWHMs are adjusted by 1.33 to reflex the magnification from the primary objective (water) to the secondary objective (air).

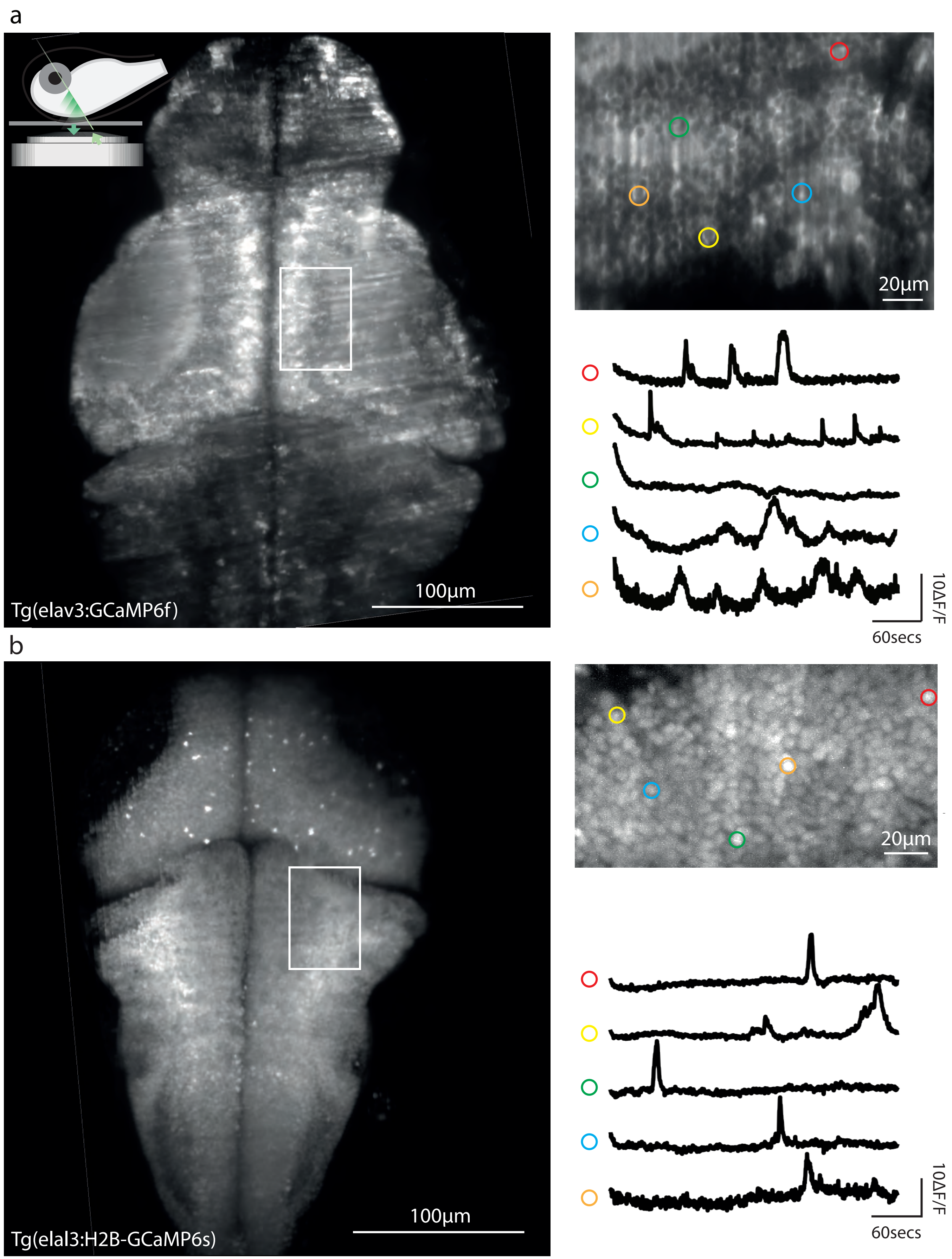


Figure 10: Whole-brain, neuron-level 3D imaging in larval zebrafish in vivo. High-resolution images are recorded in steps of 8 µm with an exposure time of 8 ms. A volume of 500 µm × 300 µm × 200 µm, containing the entire brain, is recorded once every 0.3 s. The recording of two different zebrafish lines are show in **(a)**-**(c)** and **(d)**-**(f)**. **(a)** and **(d)** show the max intensity projection of the 3D volumes. **(b)** and **(e)** show regions of interest from a single slice of the 3D stack, indicated by the write rectangles in **(a)** and **(d)**. **(c)** and **(f)** give representative fluorescence signal of five different neurons over time.

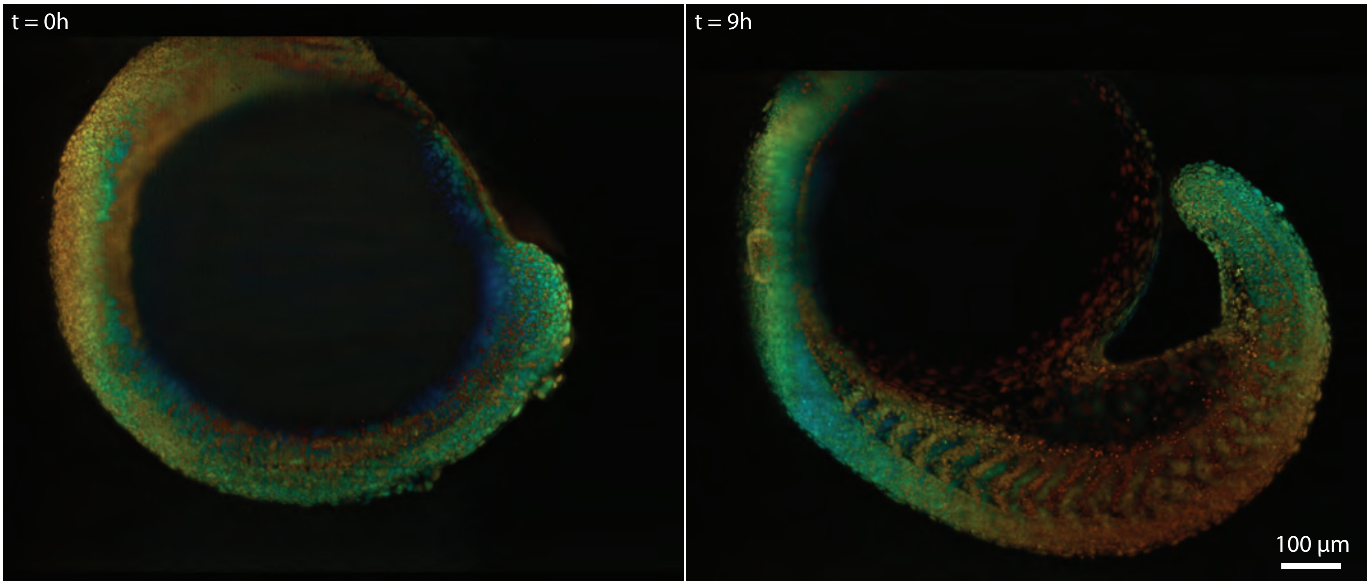


Figure 11: Volumetric imaging of zebrafish embryo development. Imaging volume is $900\,\mu\mathrm{m}\times 1000\,\mu\mathrm{m}\times 300\,\mu\mathrm{m}$ acquired every 30 seconds (two views) for 9 hours. The images shown are max-intensity projection where the depth is color-coded (blue (red) means close to (away from) the surface). See also Movie **.

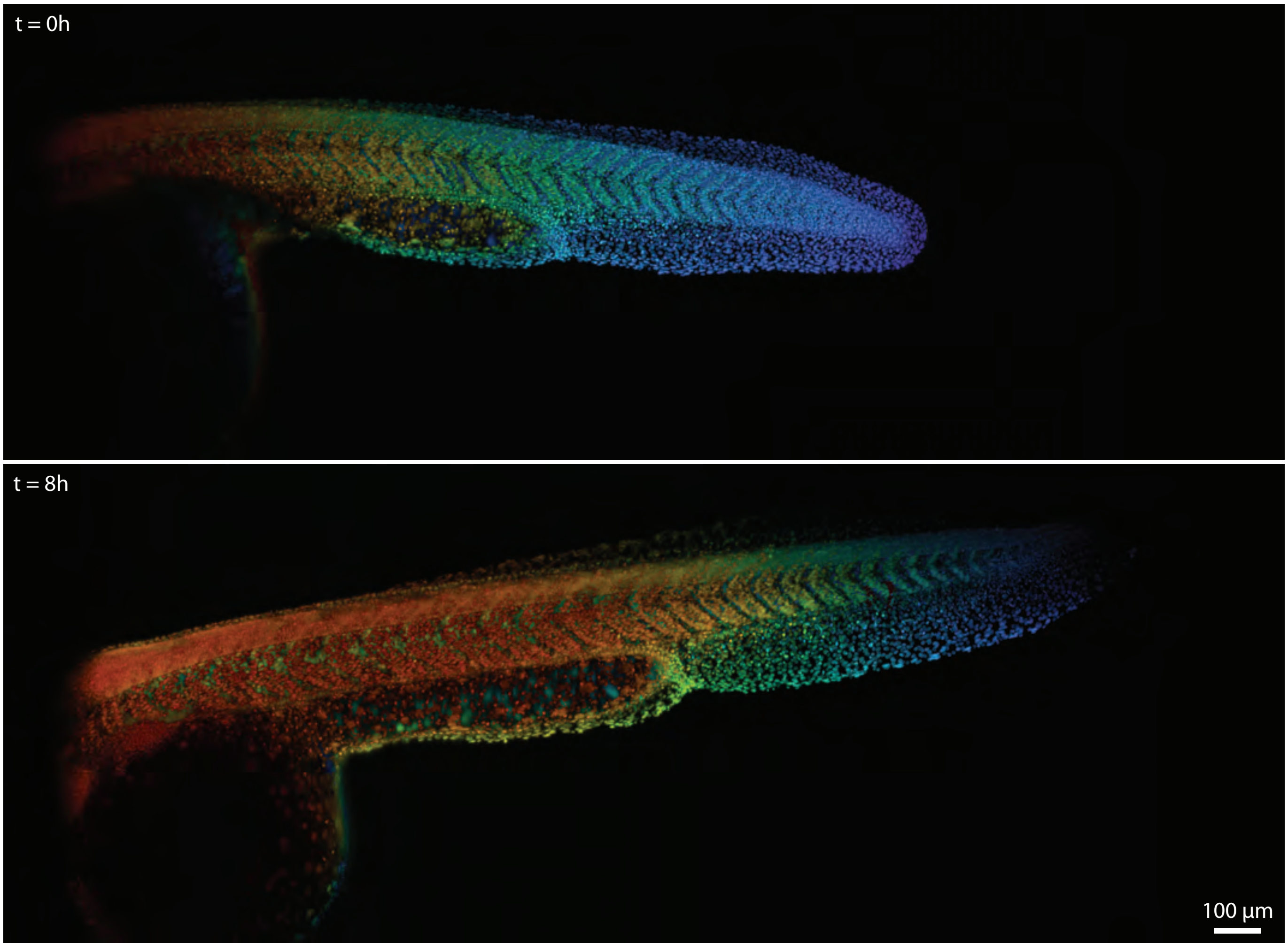


Figure 12: Volumetric imaging of zebrafish tail development. Imaging volume is 900 µm×2200 µm×300 µm acquired every 37 seconds (two views) for 8 hours. The images shown are max-intensity projection where the depth is color-coded (blue (red) means close to (away from) the surface). See also Movie **.

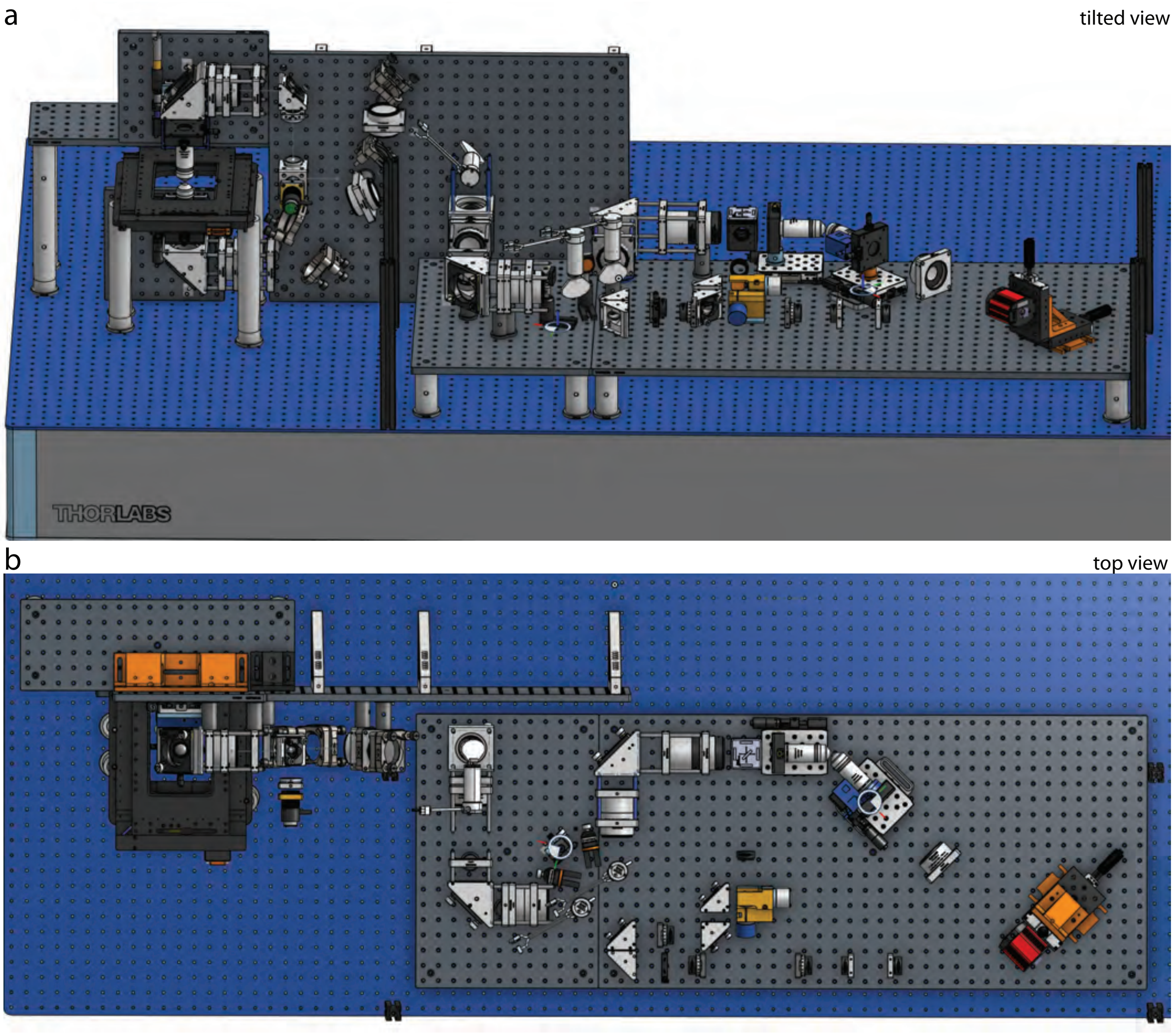


Figure 13: 3D rendering of the optical setup of the microscope. The setup is shown from two different views.

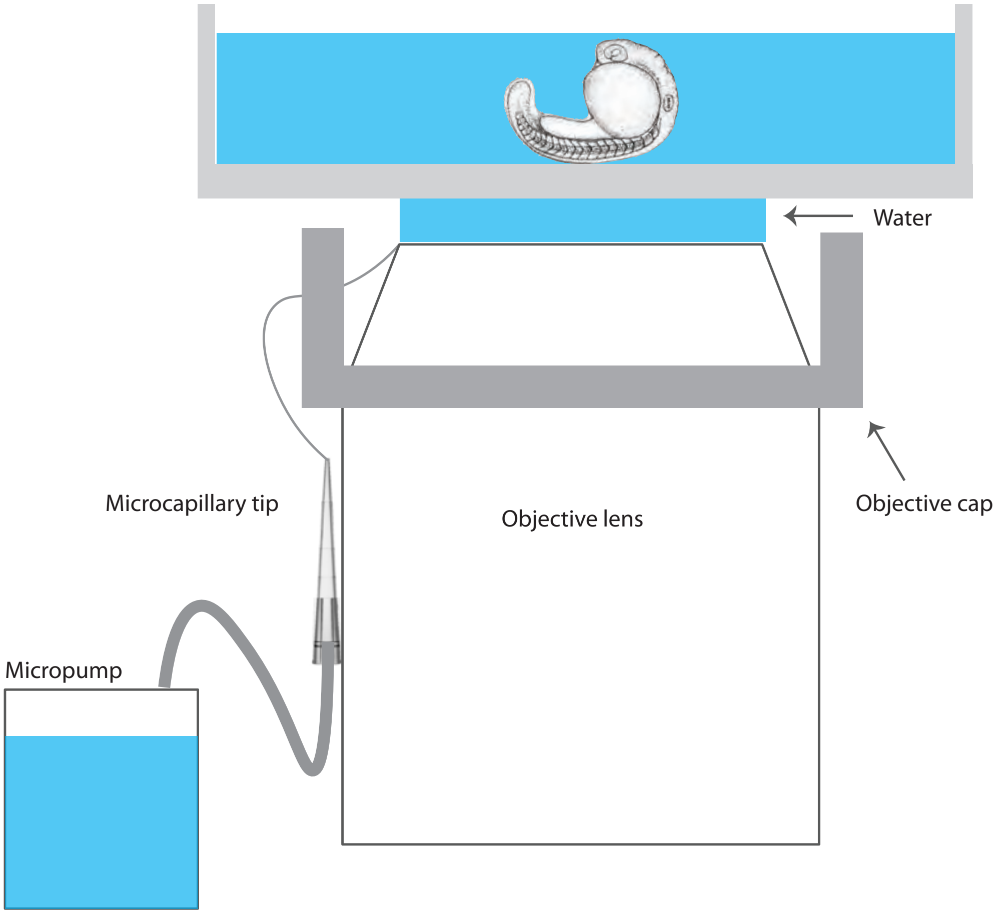


Figure 14: Water dispenser for water immersion lens. For long-term imaging, we built a water dispenser to automatically supply immersion water between the primary objective and the sample. A micropump (part of a Leica Water Immersion Micro Dispenser) pumps water through a microcapillary tip (Eppendorf Microloader) to supply the immersion water to the primary objective. A custom-designed objective cap is 3D-printed using an elastic resin (Elastic 50A, Formlabs). The cap on the objective serves as water reservoir in case of excessive water supply from the pump. The microcapillary tip is glued on the cap for support. The micropump is controlled via serial command to supply water at desired time intervals during imaging.

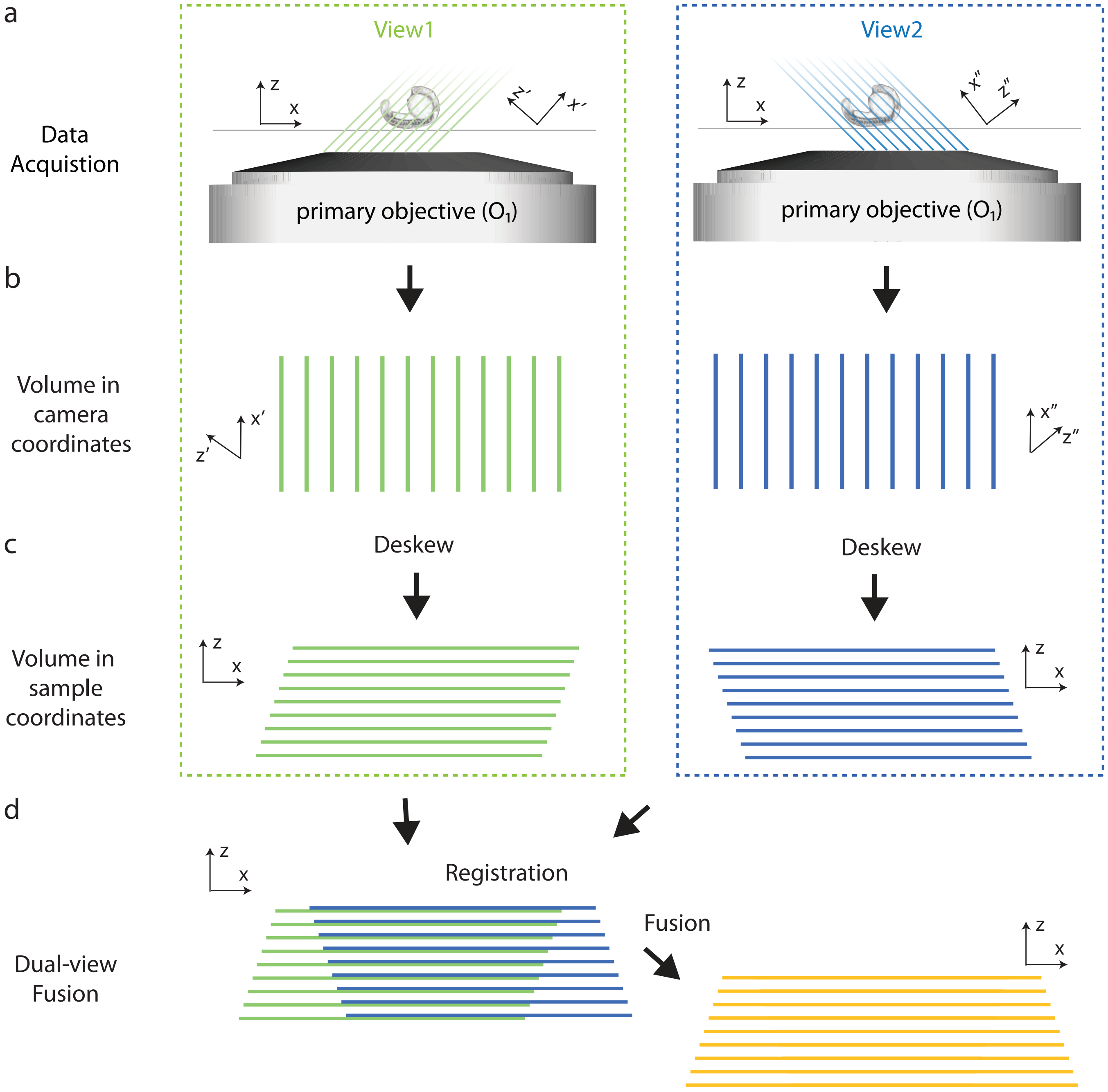


Figure 15: Data processing pipeline. The 3D stacks from each view are initially in the x'yz' and x"yz" coordinates respectively. The stacks are then resampling to the sample (i.e. xyz) coordinates. This process involves resampling of the voxels from the x'yz' (or x"yz") coordinates to the xyz coordinates, see Supp. Fig. 16. The two stacks are then registered (details) and fused (details) to a single stack.

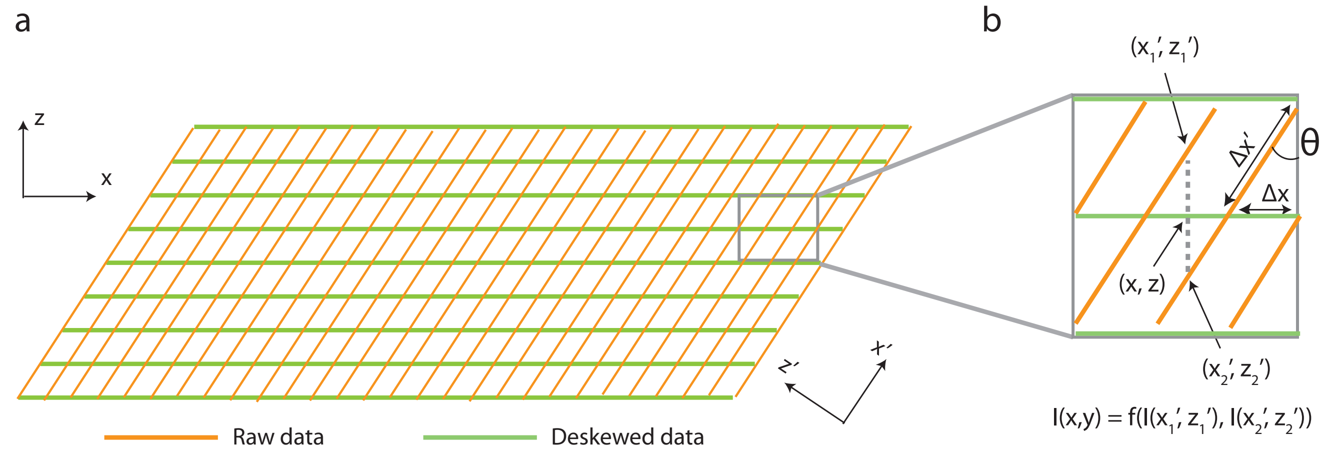


Figure 16: Resampling of data to sample coordinates. The 3D stack is initially in the x'yz' coordinates. To convert the stack to sample (i.e. xyz) coordinates, the intensity of each voxel in the xyz coordinates is the result of interpolation between two neighbouring voxels (in the x'yz' coordinates) along the vertical direction. The scanning distance between two adjacent planes in the raw data is set experimentally such that $\Delta x'$ is always an integer value of the pixel size long the x'-axis so that no interpolation is needed within along the x'-axis.

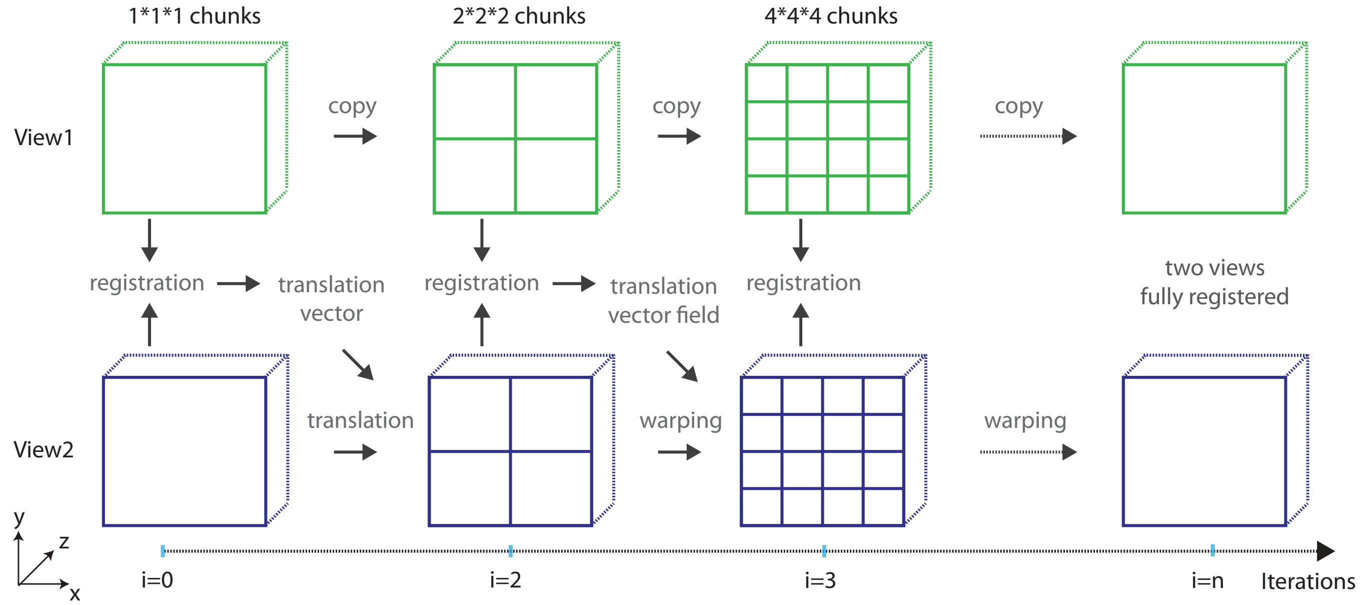


Figure 17: Warp registration. The image from the second view is registered with respective to that of the first view using an iterative multi-scale wrap approach. Within each iteration: the images are divided into chunks along all three axes by a factor of $2^i$ where i is the current iteration number; corresponding chunks from both views are registered separately with a translation model to produce a translation vector; a vector field is then calculated based on all the translation vectors; the image of the second view is warped according to the vector field. This procedure repeats until the max number of iteration or the minimal size of the chunk is reached. In this work, the max iteration number was set to 4 and the minimal chunk size 32 * 32 * 32.

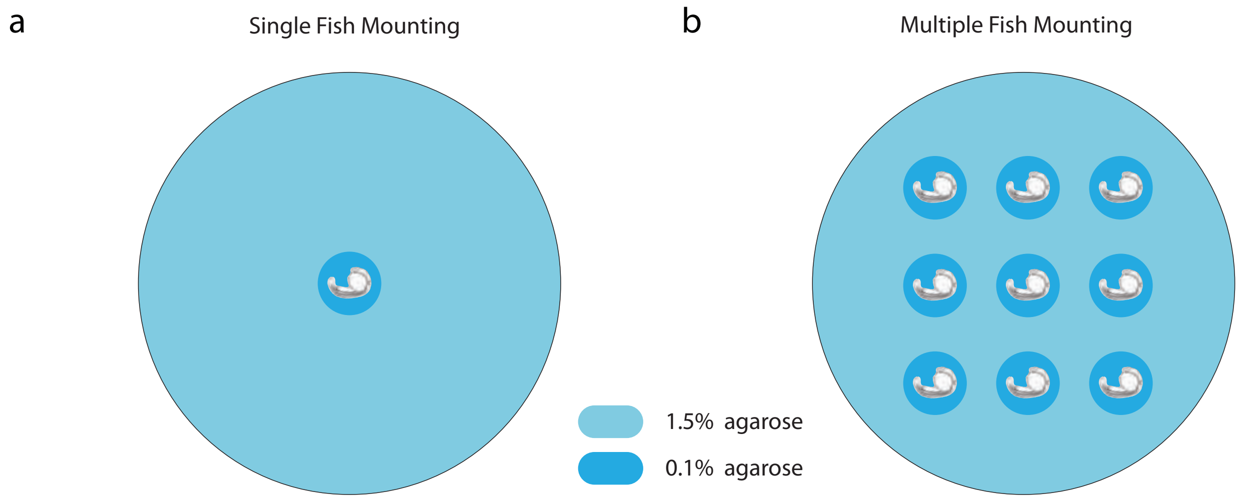


Figure 18: Zebrafish embryo mounting. 35mm cell culture dish with gass bottom (TCD35GB20, Stellar Scientific) are firstly filled with 1.5% agarose gel, with one or multiple empty spaces ($\sim$ 3mm diameter). The zebrafish embryos are then embedded in 0.1% agarose gel and mounted inside the holes.
